# Novel method for collecting hippocampal interstitial fluid extracellular vesicles (EV^ISF^) reveals sex-dependent changes in microglial EV proteome in response to Aβ pathology

**DOI:** 10.1101/2023.03.10.532133

**Authors:** Morgan C. Pait, Sarah D. Kaye, Yixin Su, Ashish Kumar, Sangeeta Singh, Stephen C. Gironda, Sami Vincent, Maria Anwar, Caitlin M. Carroll, J. Andy Snipes, Jingyun Lee, Cristina M. Furdui, Gagan Deep, Shannon L. Macauley

## Abstract

Brain-derived extracellular vesicles (EVs) play an active role in Alzheimer’s disease (AD), relaying important physiological information about their host tissues. Circulating EVs are protected from degradation, making them attractive AD biomarkers. However, it is unclear how circulating EVs relate to EVs isolated from disease-vulnerable brain regions. We developed a novel method for collecting EVs from the hippocampal interstitial fluid (ISF) of live mice. EVs (EV^ISF^) were isolated via ultracentrifugation and characterized by nanoparticle tracking analysis, immunogold labeling, and flow cytometry. Mass spectrometry and proteomic analyses were performed on EV^ISF^ cargo. EV^ISF^ were 40-150 nm in size and expressed CD63, CD9, and CD81. Using a model of cerebral amyloidosis (e.g. *APPswe,PSEN1dE9* mice), we found protein concentration increased but protein diversity decreased with Aβ deposition. Genotype, age, and Aβ deposition modulated proteostasis- and immunometabolic-related pathways. Changes in the microglial EV^ISF^ proteome were sexually dimorphic and associated with a differential response of plaque associated microglia. We found that female APP/PS1 mice have more amyloid plaques, less plaque associated microglia, and a less robust- and diverse-EV_ISF_ microglial proteome. Thus, in vivo microdialysis is a novel technique for collecting EV^ISF^ and offers a unique opportunity to explore the role of EVs in AD.

**Graphical Abstract:** Hippocampal EV^ISF^ response to amyloid beta (Aβ) is sexually dimorphic and related to the microglial EV^ISF^ proteome.

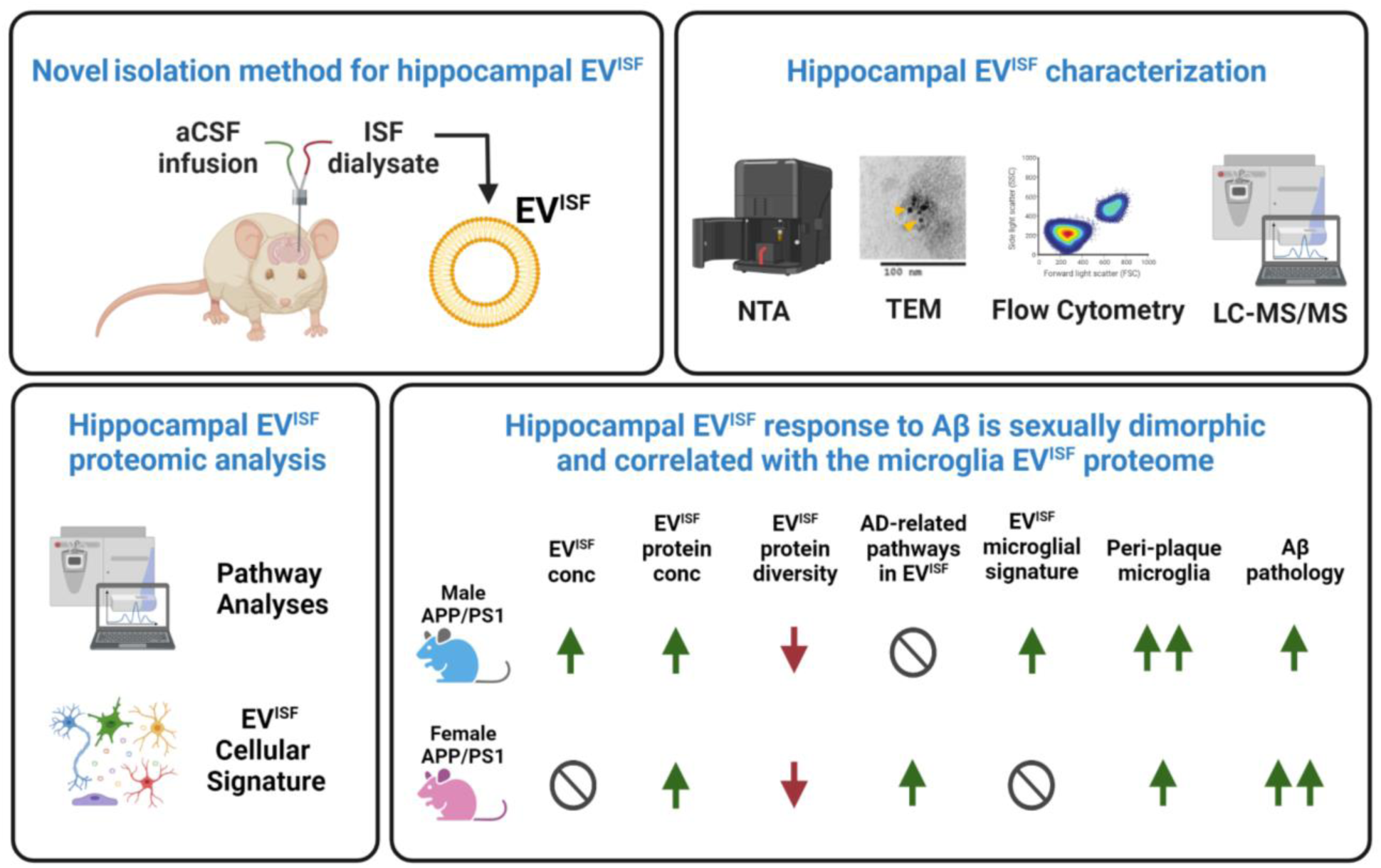

## Introduction

Brain-derived extracellular vesicles (EVs) can act as mediators and messengers of pathological conditions in Alzheimer’s disease (AD) and other neurodegenerative disorders. Small EVs (sEVs) are present in nearly all tissues and biofluids, including the brain’s interstitial fluid (ISF), cerebrospinal fluid (CSF), and blood. Exosomes are a subset of sEVs that are typically 40-150 nm in size (1). They are endosomally-derived and begin as intraluminal vesicles (ILVs) inside multivesicular bodies (MVBs) in the cell (1–3). The MVBs can either fuse with lysosomes for degradation or with the plasma membrane to release ILVs into the extracellular space as exosomes (1–4). The cargo found in exosomes and other sEVs are unique to their cell of origin and protected from degradation by a lipid bilayer as the vesicles move throughout the circulation (5,6). Their unique cargo of RNAs, proteins, and metabolites relays important physiological and metabolic information about the host cell or tissue in both homeostatic and pathological conditions. Since tissues like the brain are difficult to biopsy and brain-derived EVs can be detected in blood or CSF, EVs are attractive biomarkers to facilitate the understanding, diagnosis, and staging of AD.

While EVs can carry proteins such as amyloid-β (Aβ) and tau, the pathological hallmarks of AD, they can also relay information about the health and disease of the host cell or tissue based on their proteome, miRNAs, and metabolites (7–12). Given this, uncovering how EVs are involved in AD pathogenesis is of great interest to the field. Currently, EVs are isolated from the blood, CSF, or postmortem brain tissue from AD human patients, mouse models of AD-like pathology, and non-human primates (13,14). However, each compartment yields disparate results and has unique limitations. For example, neuronal-derived EVs in humans make up approximately 49% of brain tissue-EVs (15) but constitute only about 16% of CSF-EVs (16) and less than 15% of plasma-EVs (17). Microglia/macrophage-derived EVs make up only 4-12% of CSF-EVs, which further differs from estimations in the blood (18). Similarly, miRNAs from brain-versus plasma-derived EVs from AD patients often fail to correlate with one another (19). This suggests that compartments distal to the brain do not always reveal all disease-related or cell type-specific changes in brain EV populations, potentially biasing our understanding of disease progression. Brain-derived EVs isolated from CSF and blood also represent a heterogeneous pool of EVs and could fail to reveal brain region-specific changes associated with disease progression.

Over the past decade, advanced protocols for isolating sEVs from the extracellular space of postmortem brain tissue were developed to understand region-specific changes associated with pathology (20–29). Implementation of these methods demonstrated that levels of sEVs are differentially modulated by various AD risk factors (e.g. ApoE4, Down syndrome, APP) as a function of age and disease severity (23, 30-34). There are several limitations with this approach. First, EVs from postmortem brains may not fully reveal the active role EVs play in the AD cascade since they are sampled at one time-point, typically at an end stage of the disease, following a highly variable or lengthy postmortem interval. Thus, any dynamic changes in the in vivo brain could be lost with this approach. Some of these methods use harsh methods of EV extraction, disrupting the brain cells which potentially leads to contamination of EVs with ILVs and other intracellular vesicles. Further, many of these studies are performed on whole brain, hemibrain, or pooled samples, losing the region-specific changes that occur in AD. Thus, despite these advances in brain EV isolation methods, the problem of collecting EVs from the extracellular space of an intact, in vivo brain, in a region-specific manner, persists. Therefore, developing new methods that detect dynamic changes in the EV pool in the brain’s ISF in live animal models of AD is imperative.

Isolating EVs from the live brain can help us elucidate the role EVs play in AD-pathogenesis as well as identify novel biomarkers of AD pathogenesis. Thus, we developed a novel method for collecting EVs from the hippocampal ISF of unanesthetized, freely-moving mice using in vivo microdialysis. To date, in vivo microdialysis is widely utilized in animal models of AD-like pathology for collecting Aβ, tau, and other proteins and metabolites but heretofore untested as a method for collecting EVs from the ISF (35–44). By coupling in vivo microdialysis and EV isolation with proteomic profiling and bioinformatics, we hypothesized that this approach would allow us to characterize brain region-and cell type-specific changes in EVs relative to Alzheimer’s-like pathology in vivo. In this study, we examined how the hippocampal ISF population of sEVs changes in a mouse model of Alzheimer’s-related pathology, APPswe, PSEN1dE9 (APP/PS1).

We found that in vivo microdialysis successfully captures sEVs from the hippocampal ISF (EV^ISF^) of live mice. Ultracentrifugation (UC) was the preferred method of isolating EV^ISF^ compared to ExoQuick and Size Exclusion Chromatography (SEC). Interestingly, age, sex, and amyloid pathology modulated EV^ISF^ characteristics, specifically EV^ISF^ number. Proteomic and bioinformatic analysis of EV^ISF^ revealed that this method of EV collection selects for a unique EV population disparate from other studies. Surprisingly, EV^ISF^ protein concentration increased while protein diversity within the EV^ISF^ decreased with Aβ pathology. EV^ISF^ also revealed changes in proteostasis and neuroinflammation as early as 3-months in APP/PS1 vs WT mice. Aβ deposition further perturbed proteostasis and inflammation-related contents of EV^ISF^. Age, Aβ deposition, and sex differentially modulated the EV^ISF^ proteome based on the cell type of origin. Specifically, microglia-derived proteins correlated with ISF Aβ, Aβ deposition, and markers of microglial activation, but only in EV^ISF^ of male APP/PS1 mice. We also observed sex dependent changes in plaque associated microglia that could be related to changes in the EV^ISF^ microglial proteome. Thus, this study demonstrates for the first time that not only can EV^ISF^ be collected from an in vivo brain using microdialysis, but this population of EVs is modulated by genotype, age, and Aβ pathogenesis as well. Finally, EV^ISF^ can detect sex-dependent changes in microglia-related EV proteins that correlate with AD-related pathology.

## Methods

### Animals

3-and 9-month-old male (n=46) and female (n=19) mice heterozygous for the APPswe,PSEN1dE9 mutation (45) (APP/PS1) or littermate controls (WT) on a B6C3 background were used in this study. Mice were given food and water ad libitum and maintained on a 12:12 light/dark cycle (46–48). All protocols were approved by the Institutional Animal Care and Use Committee at Wake Forest School of Medicine.

### Workflow of EV^ISF^ collection, isolation, and analysis

Hippocampal ISF was collected from 3-and 9-month-old APP/PS1 and WT mice using in vivo 1000kDa microdialysis. Artificial CSF (aCSF) was infused at 1.2 µL/min with a syringe pump. ISF was collected via a peristaltic pump at 1 µL/min into a 4°C refrigerated fraction collector. ExoQuick, Size Exclusion Chromatography (SEC), and Ultracentrifugation (UC) were tested in order to determine the preferred method of EV^ISF^ isolation. Ultimately, EV^ISF^ were isolated via UC and characterized by nanoparticle tracking analysis (NTA), transmission electron microscopy (TEM), and flow cytometry. Liquid chromatography-mass spectrometry/mass spectrometry (LC-MS/MS) was performed on the EV^ISF^ protein cargo. Proteomic analyses were carried out by utilizing the IPA, DAVID, and STRING databases. The cell type sources of EV^ISF^ proteins were determined using the publicly available brainrnaseq.org database (49). The percentages of cell type-related proteins in EV^ISF^ were revealed. Additionally, postmortem analyses were performed on the ISF and brain tissue via enzyme-linked immunosorbent assays (ELISAs) and immunohistochemistry (IHC), respectively (Figure 1).

**Figure 1:**
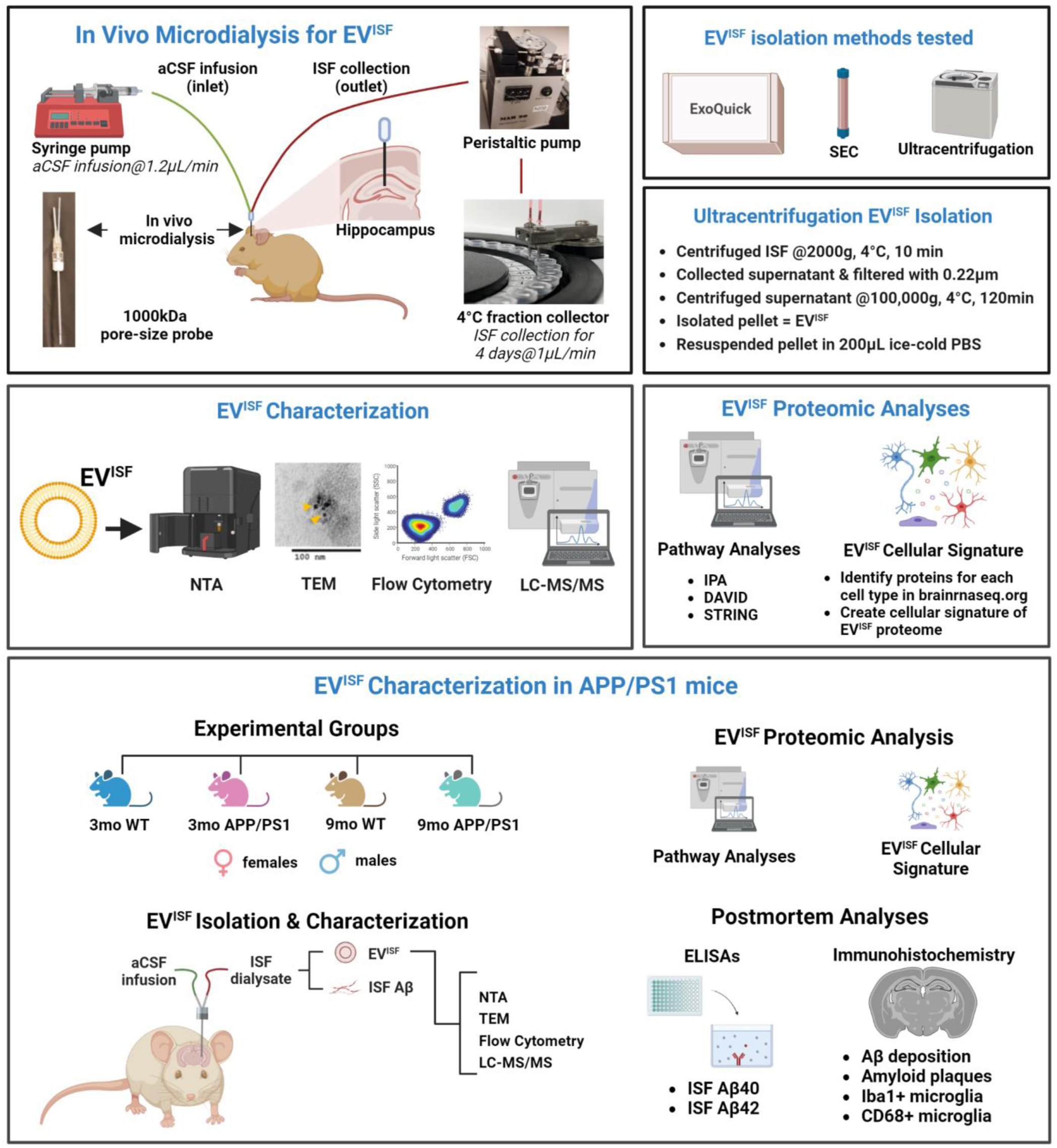
Technical development of hippocampal interstitial fluid (ISF) extracellular vesicles (EV^ISF^) isolation, characterization, and analysis. Workflow of collection, isolation, characterization, and proteomic analyzation of EV^ISF^. (*In Vivo Microdialysis for EV^ISF^*) ISF was collected from the hippocampus via in vivo microdialysis utilizing a 1000kDa pore-size probe. (*EV^ISF^ isolation methods tested*) Three different methods for EV isolation (ExoQuick, Size-Exclusion Chromatography, and Ultracentrifugation) were tested on ISF. (*Ultracentrifugation EV^ISF^ isolation*) Steps for isolating EVs from the ISF using ultracentrifugation. (*EV^ISF^ Characterization*) EVs were characterized by nanoparticle tracking analysis (NTA), transmission electron microscopy (TEM), flow cytometry, and LC-MS/MS. *(EV^ISF^ Proteomic Analyses)* Pathway analyses (IPA, DAVID, and STRING) were performed on the EV^ISF^. The EV^ISF^ cellular signature was determined via utilization of the brainrnaseq.org database (49) to identify the cell-type origins of the EV^ISF^ proteins. *(EV^ISF^ Characterization in APP/PS1 mice)* EV^ISF^ from the hippocampus of 3mo and 9mo WT and APP/PS1 mice, both sexes, were isolated and characterized. Proteomic analyses were performed on the EV^ISF^. Finally, postmortem analyses were performed on the collected ISF and brain tissue.

### In vivo 38kDa microdialysis

In vivo microdialysis was performed on mixed 3-month-old APP/PS1 and WT male and female mice (n=6) as previously described (42, 43, 50). Briefly, a guide cannula (BR-style, BASi Instruments) was stereotaxically implanted into the hippocampus (from bregma, A/P: −3.1 mm; M/L: −2.5 mm; D/V: −1.2 mm; at 12° angle) and secured into place with dental cement. Mice were transferred to Raturn recording chambers (BASi Instruments) then allowed to recover for ∼72 hrs. A microdialysis probe (2 mm; 38kDa molecular weight cut off; BR-style; BASi) was inserted into the guide cannula then connected to a syringe pump (KD Scientific). 0.15% bovine serum albumin (BSA, Sigma) in artificial cerebrospinal fluid (aCSF: 1.3 mM CaCl2, 1.2 mM MgSO4, 3 mM KCl, 0.4 mM KH2PO4, 25 mM NaHCO3 and 122 mM NaCl; pH = 7.35) was filtered with a 0.10µM membrane and infused using the syringe pump (KD Scientific) at a flowrate of 1 μL/min. ISF was collected at a flowrate of 1 μL/min every 90 min into polypropylene tubes in a refrigerated fraction collector (MAB 85, SciPro) for ∼72 hrs. The first 6 hrs of collection were excluded during acclimation. The remaining fractions were pooled into 15mL polypropylene tubes yielding a total volume of ∼4 mL. Pooled samples were stored at −80°C as there was no difference between EVs isolated from fresh versus frozen ISF. Following in vivo microdialysis, mice were euthanized via transcardial perfusion with cold Dulbecco’s phosphate buffered saline (dPBS; gibco Life Technologies Corporation, NY) with 0.3% heparin. The brain was bisected for biochemical and IHC analyses.

### In vivo 1000kDa microdialysis

In vivo 1000kDa microdialysis was performed on a separate cohort of 3-month WT males (n=21) for EV isolation methods and another cohort of 3-and 9-month-old APP/PS1 and WT mice (n=4-6 mice/sex/age/group) for EV characterization and LC-MS/MS of EV^ISF^. A modified version of previously published methods for in vivo microdialysis was used (41, 46, 51, 52). A guide cannula (AtmosLM PEG-12, Amuza, Eicom) was stereotaxically implanted into the left hippocampus (from bregma, A/P: −3.1 mm, M/L: −2.5 mm, D/V: −1.2 mm at 12° angle) and secured into place with dental cement. Mice were transferred to Raturn recording chambers (BASi) then allowed to recover for ∼72 hrs. After the recovery period, a microdialysis probe (1000kDa probe: 1000kDa cut off, AtmosLM PEP-12-02, Amuza, Eicom) was inserted into the guide cannula. 0.15% bovine serum albumin (BSA, Sigma) in artificial cerebrospinal fluid (aCSF: 1.3 mM CaCl2, 1.2 mM MgSO4, 3 mM KCl, 0.4 mM KH2PO4, 25 mM NaHCO3 and 122 mM NaCl; pH = 7.35) was filtered with a 0.10µM membrane and infused using a syringe pump (KD Scientific) at a flowrate of 1.2 μL/min. ISF was collected at a flowrate of 1 μL/min via push-pull method with a peristaltic pump (MAB 20, SciPro). Samples were collected every 90 min into polypropylene tubes in a refrigerated fraction collector (MAB 85, SciPro) for ∼72 hrs. Fractions were pooled, excluding first 6 hrs of acclimation, into 15mL polypropylene tubes yielding a total volume of ∼4 mL. Pooled samples were stored at −80°C as no difference was determined between analyzation of EVs isolated from fresh versus frozen ISF. Following in vivo microdialysis, mice were euthanized via transcardial perfusion with cold dPBS (gibco Life Technologies Corporation, NY) with 0.3% heparin. The brain was bisected for biochemical and IHC analyses.

### Total EV isolation

Total EVs (TE) were isolated from ISF samples using three methods: **(1) Ultracentrifugation (UC):** ISF was harvested and centrifuged at 2000 *g* at 4°C for 10 minutes to remove cell debris. The supernatant was collected and filtered through 0.22 µm filters to remove large sized vesicles and further centrifuged (Beckman Coulter Optima Ultracentrifuge, Beckman Coulter, Brea, CA) at 100,000 *g* at 4°C for 120 minutes with a Type 70 Ti rotor to pellet sEVs. The supernatant was carefully removed, and sEV-containing pellets were resuspended in 200µL of ice-cold PBS. **(2) ExoQuick precipitation:** ISF samples were centrifuged at 3000 × g for 15 minutes to remove cells and cell debris. The supernatant was collected in a sterile tube, and an appropriate volume of ExoQuick-TC was added, mixed, and incubated overnight at 4^°^C before centrifugation at 1500 × g for 30 minutes. The supernatant was collected and discarded, and the pellet was resuspended in sterile 1X PBS. **(3) Size exclusion chromatography (SEC):** EVs were isolated from ISF using Sephadex-G25 (GE Healthcare; Uppsala, Sweden) following the vendor’s instruction. The column was washed with PBS (pH 7.4, 0.22 µm filtered). Subsequently, the column was equilibrated with 25 mL PBS and sample was added to enter the gel bed completely. The purified sample was eluted with an appropriate volume of PBS. The eluate was collected in 10 sequential fractions of 1 mL and each fraction was analyzed for particle concentration (number/mL) by nanoparticle tracking analysis (NTA) as described below. Protein concentration of EVs was measured by a NanoDrop.

### Nanoparticle tracking analyses (NTA)

Quantification of the hydrodynamic diameter distribution and concentration (number/mL) of EV were performed using the Nanosight NS300 (Malvern Instruments, UK) equipped with a violet laser (405 nm) and running software version NTA3.4. The instrument was primed using phosphate-buffered saline (PBS), pH 7.4 and the temperature was maintained at 25°C. Five measurements (30 seconds each) were obtained for each sample, and their average was plotted as representation of size distribution and concentration (particles/ml).

### Flow cytometry

Flow cytometry analysis was performed to evaluate the percentage of typical EV tetraspanin markers following methods reported recently (53). TE were labeled with membrane labeling dye CellBrite 488 (Biotium, California, USA) with and without the CD63-APC and CD81-PE (both from BioLegend, CA, USA) antibodies. EVs without dye were used as control to set the gate for positively labeled EVs. EVs labeled with dye but without CD63-APC/ CD81-PE antibody were used to set the gate for APC/ PE positive events. CD63-APC/ CD81-PE antibody and dye at the same dilution in PBS (filtered through 0.22-micron filter) were also analyzed. All samples were acquired on CytoFlex (Beckman Coulter Life Science, Indianapolis, United States) for 60 seconds at a low flow rate. Filtered PBS was run for 60 seconds in between the samples. Following sample acquisition, EVs were lysed by adding 0.25% triton-100 and acquired again to confirm the captured events were EVs.

### Immunogold labeling

EVs were fixed with 2% paraformaldehyde in PBS buffer (pH 7.4), then adsorbed for 1 hour to a carbon-coated grid. EVs were incubated with CD63 (Abcam, Cat. No. ab59479) or CD9 (Thermo Fisher, Cat. No. MA5-31980) primary antibody. A secondary antibody tagged with 10 nm gold particles was used. EVs were contrasted in 1% uranyl acetate for 5 minutes, and images were captured on Tecnai T12 transmission electron microscope (TEM). Samples treated with only gold-labeled secondary antibody were used as a negative control.

### EV^ISF^ Proteomic Profiling via LC-MS/MS

EV^ISF^ were isolated from ISF by ultracentrifugation as described above. EV^ISF^ were briefly treated with trypsin at 1:100 (enzyme:EV^ISF^ protein) ratio for 2 hrs at 37°C to remove corona or surface proteins, and shaved EVs were pelleted again by ultracentrifugation at 200,000 x g for 3 hrs. Core proteins were prepared for LC-MS/MS analysis according to the method reported by us recently (14,54). Samples were analyzed on a LC-MS/MS system consisting of an Orbitrap Velos Pro Mass Spectrometer (Thermo Scientific, Waltham, MA) and a Dionex Ultimate3000 nano-UPLC system (Thermo Scientific, Waltham, MA). An Acclaim PepMap 100 (C18, 5 μm, 100 Å, 100 μm x 2 cm) trap column and an Acclaim PepMap RSLC (C18, 2 μm, 100 Å, 75 μm x 50 cm) analytical column were employed for peptide separation. MS spectra were acquired by data-dependent scans, and then searched using Sequest HT within the Proteome Discoverer v2.2 (Thermo Scientific, Waltham, MA) and UniProt human protein FASTA database (20,258 annotated entries, Feb 2018). Search parameters were as follows: FT-trap instrument, parent mass error tolerance of 10 ppm, fragment mass error tolerance of 0.6 Da (monoisotopic), variable modifications of 16 Da (oxidation) on methionine. Fixed modification of 57 Da (carbamidomethylation) was applied to cysteine for core protein identification.

### Determination of cell type-related proteins in EV^ISF^

The gene names of EV^ISF^ proteins were determined using UniProt.org (55). Brainrnaseq.org database (49) determined cell type origin(s) (microglia, neuron, oligodendrocyte, astrocyte, endothelial cell) of EV^ISF^ proteins. If a protein was represented in more than one cell type, it was counted for each cell type it was found in. Within each animal, the total number of times a particular cell type was identified was summed. A percentage was determined by dividing that summed number by the sum total of all cell types found in that animal and multiplying by 100. Significant effects were determined using one-way ANOVAs with Tukey’s multiple comparisons post-hoc tests.

### Immunohistochemistry and quantification for Aβ and CD68 staining

3-month and 9-month WT and APP/PS1 mice were anesthetized with isoflurane and transcardially perfused with cold dPBS with 0.3% heparin. The brains were removed and fixed in 4% paraformaldehyde for at least 48 hrs at 4°C. Prior to sectioning, the brains were cryoprotected in 30% sucrose then sectioned on a freezing microtome at 50μm. Three serial sections (300μm apart) through the anterior-posterior aspect of the hippocampus were immunostained for either Aβ deposition or CD68. <u>Aβ deposition.</u> Free-floating sections were stained for Aβ deposition using a biotinylated, HJ3.4 antibody (anti-Aβ_1–13_, mouse monoclonal antibody, a generous gift from the Holtzman Lab, Washington University). <u>CD68 staining.</u> Free floating sections were blocked with 3% donkey serum and incubated overnight with rat anti-CD68 (1:200, Bio-Rad). Aβ deposition and CD68 staining were both developed with a Vectastain ABC kit and DAB reaction. The brain sections were imaged with the Wake Forest Imaging Core NanoZoomer slide scanner and quantified as described previously (37,46). Data is represented as area fraction means ±SEM. Mean reactivity and percent area occupied by HJ3.4 or CD68 were quantified with ImageJ software (National Institutes of Health) by a blinded researcher as previously described (37,46). Statistical significance was determined using a one-way ANOVA with multiple comparisons Tukey’s tests.

### Immunohistochemistry and quantification for X34, IBA1, and CD68 staining

Three 50_μm brain sections of the anterior cortex were taken from 3-month-old and 9-month-old WT and APP/PS1 mice. Free-floating sections were incubated in X34 (Sigma) for 20 minutes at room temperature in a 60%_PBS/40%_EtOH mix. Sections were washed briefly three times with 60%_PBS/40%_EtOH and then two times in PBS before blocking. Following blocking, sections were incubated in IBA1 (Wako; anti-rabbit 1:1000) and CD68 (Bio-Rad; anti-mouse 1:150) and kept at 4_°C overnight with slow agitation. The following morning, sections were washed with PBS and then blocked and incubated with corresponding secondaries for 2 hours at room temperature. The secondary for IBA1 was goat-anti-rabbit Alexa fluorophore 488 (Invitrogen/Thermo Fisher) while the secondary for CD68 was goat-anti-mouse Alexa fluorophore 594 (Invitrogen/Thermo Fisher). After the 2-hour incubation, sections were briefly washed in PBS again and mounted with fluoromount and coverslipped. Sections were imaged using a Nikon Ci laboratory microscope. For all analyses, images were acquired using a 20x objective with 2880×2048 pixel resolution. For quantification of plaques, microglia, and CD68, images were imported to Fiji and thresholding methods were performed for each individual channel. After thresholding, the area fraction was identified for each channel and each image. Mixed-effects analyses were performed and followed by Tukey’s post hoc tests to identify group differences. In addition to area fraction measurements, all three channels for each image were merged to perform colocalization analyses using EzColocalization (56). A threshold overlap score (TOS), a measure of colocalization, was determined for each image and compiled for each group. Differences in TOS between the groups were assessed using a mixed-effects analysis and Tukey’s post-hoc analyses.

### *A*β*40 and A*β*42 ELISAs*

Hippocampal ISF samples from 3-month and 9-month APP/PS1 mice and WT mice (n=4-6/group) collected from microdialysis experiments were analyzed for Aβ40 and Aβ42 using sandwich ELISAs as previously described (35, 37, 39, 40). Briefly, Aβ40 & Αβ42 were quantified using monoclonal capture antibodies (generous gifts from Dr. David Holtzman, Washington University) targeted against amino acids 33-40 (HJ2) or 35-42 (HJ7.4) respectively. For detection, a biotinylated monoclonal antibody against the central domain amino acids 13-28 (HJ5.1B) was used, followed by streptavidin-poly-HRP-40 (Fitzgerald). The assay was developed using Super Slow TMB (Sigma) and the plates were read on a Bio-Tek Synergy H1 plate reader at 650 nm. Standard curves with known Aβ standards related optical density measures to Aβ concentrations. Statistical significance was determined using a one-way ANOVA. Data is represented by means ±SEM.

### Data analysis

<u>IPA analysis</u> The proteomics data was analyzed using Ingenuity Pathway Analysis (IPA) (QIAGEN Inc.) as previously described (64–66). Briefly, identified proteins were functionally assigned to canonical pathways and subsequently mapped to the most significant networks generated from previous publications and public protein interaction databases. Canonical pathways were presented as –log_10_(p-value). <u>DAVID analysis:</u> Database for Annotation, Visualization and Integrated Discovery (DAVID) Bioinformatics Resources 6.8 was utilized for Gene Ontology (GO) analysis of hippocampal EV^ISF^ proteins. GO terms for the top “Biological process”, “Cellular component”, “Molecular function”, and “Tissue” were presented as-log_10_(FDR p-value) ≥ 1.3. Kyoto Encyclopedia of Genes and Genomes (KEGG) pathways obtained via DAVID were also presented as –log_10_(FDR p-value) ≥ 1.3. <u>STRING analysis:</u> STRING database (57) was utilized for determining the top 20 biological processes of EV^ISF^ proteins. <u>Correlation analyses:</u> To define the relationships between all metrics measured, correlational analyses were performed between the following variables to determine the relationship: 1) ISF Aβ40, 2) ISF Aβ42, 3) Aβ deposition, 4) CD68 staining, and 5) EV^ISF^ cell type % (neuronal, astrocyte, etc.). Correlation coefficients (r), coefficients of determination (R^2^), and p-values (p) were calculated for each relationship to define the strength of the interaction. <u>Histological and biochemical analyses:</u> IHC images were evaluated for percent area of Aβ+ and CD68+ in the hippocampus and cortex using ImageJ (National Institute of Health) software and compared to WT. Additionally, all histological markers were compared between sexes to evaluate difference in AD-related burden. Significant effects were determined using one-way ANOVAs with post-hoc tests. One-way ANOVAs or t-tests compared histochemical and biochemical readouts between groups. Statistical significance for all analyses was set at p < 0.05. Prism 9 (GraphPad Software, Inc., San Diego, CA) was used for statistical analyses. Venn diagram comparing the EV^ISF^ proteins from the four mouse groups was created using R Studio R package version 1.7.1.R “VennDiagram: Generate High-Resolution Venn and Euler Plots” by Hanbo Chen (2021) https://CRAN.R-project.org/package=VennDiagram. Three-way comparison of the EV^ISF^ proteins, Choi et al EV dataset (58), and ExoCarta Top 100 list (59) was generated via the Whitehead Institute for Biomedical Research Bioinformatics and Research Computing public tools website: barc.wi.mit.edu/tools/compare_3_lists/. The three-way comparison of the EV^ISF^ protein dataset to the human brain tissue-derived (15) and mouse brain tissue-derived (60) EV datasets was also generated via barc.wi.mit.edu/tools/compare_3_lists/.

## Results

### In vivo microdialysis as a novel method for collecting EV^ISF^ from hippocampus of live, freely moving, unanesthetized mice

Two different microdialysis probes with varying pore sizes, 38kDa and 1000kDa, were used to determine the best approach for isolating EV^ISF^ from live, freely moving mice (Figure 2A). For our initial experiments, ISF was collected from the hippocampus of 3-month-old WT and APP/PS1 mice using 38kDa and 1000kDa probes. EV^ISF^ were isolated via ultracentrifugation. Although 38kDa and 1000kDa probes collected EV^ISF^ in the sEV size range (<200 nm) (61), the 1000kDa probe collected relatively larger 100-150nm sEVs compared to the 38kDa probe which collected 50-125nm sized EV^ISF^ (p=0.0003; Figure 2B). The concentration of EV^ISF^ collected with the 1000kDa probe was relatively lower when compared to the 38kDa probe (p<0.0001; Figure 2C), while the protein concentration of EV^ISF^ collected via the 1000kDa probe was higher than the 38kDa probe (p=0.0143; Figure 2D). When the EV^ISF^ protein concentration was divided by the concentration of EV^ISF^ (number /ml), the 1000kDa probe collected sEVs with greater protein concentration/sEV (p=0.0043; Figure 2E). Although the 38kDa pore size probe can collect various forms of Aβ in the ISF (39,62), a larger pore size probe like the 1000kDa is required to collect larger proteins such as ApoE (63) or tau (41). Thus, since the 1000kDa probe collected sEVs with more protein concentration, and potentially larger protein species within the sEVs, the 1000kDa probe was chosen for the development and implementation of this novel method of EV^ISF^ isolation.

**Figure 2:**
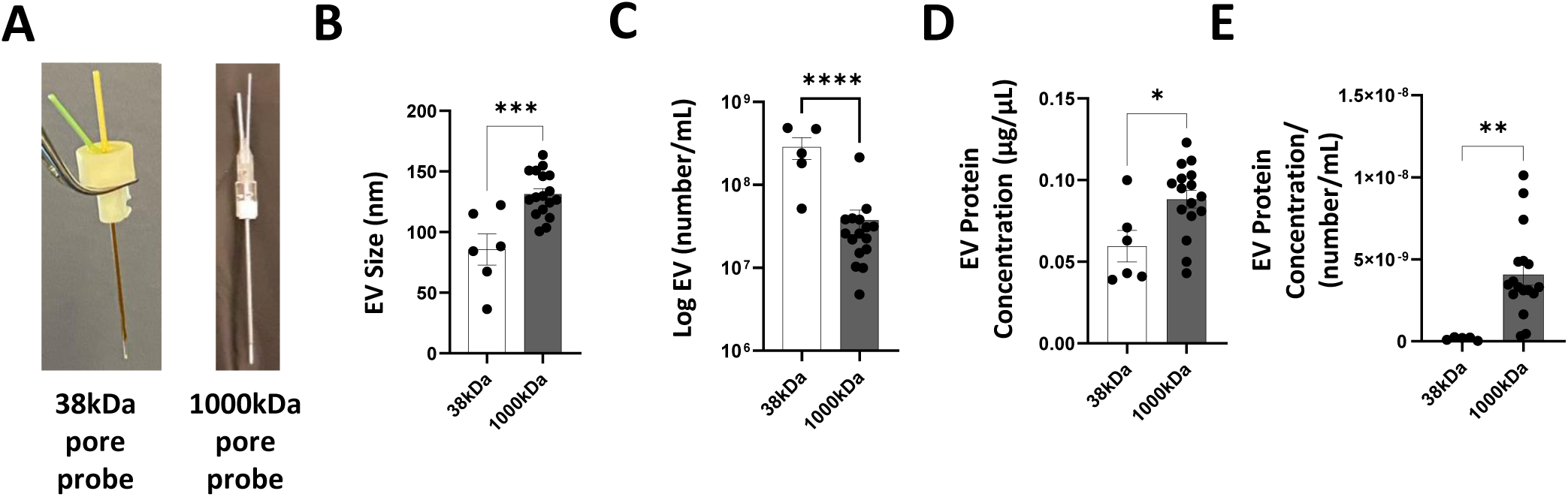
Comparing *in vivo* microdialysis methods for successful collection of hippocampal EV^ISF^. **(A)** Probe comparison: 38kDa pore size vs 1000kDa pore size for EV^ISF^ collection via *in vivo* microdialysis. Hippocampal ISF was collected from 3mo WT and APP/PS1 mice via 38kDa and 1000kDa microdialysis. EV^ISF^ was isolated via ultracentrifugation. **(B)** The mean size of EV^ISF^ collected via 1000kDa probe was larger (131nm) than those collected via 38kDa (86nm) (p=0.0003). **(C)** EV^ISF^ concentration (particle number/mL) was lower via 1000kDa collection than 38kDa because 1000kDa probes allowed collection of larger sized EV^ISF^ (p<0.0001). **(D)** However, EV^ISF^ collected via 1000kDa had greater protein concentration than 38kDa EV^ISF^ (p=0.0143). **(E)** The 1000kDa pore size probe collects EVs with greater protein concentration than the 38kDa (p=0.0043). p<0.05*, p<0.01**, p<0.001***, p<0.0001****, unpaired, two-tailed, Student’s t-test. N=5-16 per group, both sexes.

### Ultracentrifugation as the preferred method for isolating EV^ISF^

Next, we explored which isolation method - ultracentrifugation (UC), ExoQuick, or Size Exclusion Chromatography (SEC) - was best for isolating EV^ISF^. NTA was used to characterize the mean size and concentration of EV^ISF^ isolated for each method. When comparing EV size, ExoQuick isolated EV^ISF^ with a larger size compared to UC and SEC (Figure 3A, left panel). However, a relatively greater concentration of EV^ISF^ was isolated with UC versus ExoQuick or SEC (Figure 3A, right panel). When comparing the percentage of EV concentration for different EV size ranges, EV^ISF^ isolated with UC and SEC evidenced a higher percentage of EVs in the 100-150nm EV size range (Figure 3B). However, ExoQuick isolated more EV^ISF^ in the 150-200nm and >300nm ranges (Figure 3B). EV^ISF^ isolated via all three methods were analyzed via flow cytometry in order to determine the percentage of CD63-and CD81-positive EV^ISF^, markers used to identify exosome-enriched sEVs. More CD63-positive EV^ISF^ were detected via UC isolation (61.73%) compared to ExoQuick and SEC methods (48.17% and 56.14%, respectively) while SEC resulted in more CD81-positive EV^ISF^ (80.99%) compared to both ExoQuick and UC (71.38% and 67.56%, respectively) (Figures 3C-D). Finally, EV^ISF^ isolated via UC underwent immunogold labeling for CD63 and CD9, which demonstrated that tetraspanin EV markers were present on the surface of the EV^ISF^ (Figure 3E). Since UC resulted in a greater number of EV^ISF^ that also fall in the sEV size range and expressed the markers associated with an exosome-enriched EV population, UC was chosen as the preferred method of EV^ISF^ isolation. Nevertheless, ExoQuick and SEC also supported the isolation of EVs from the ISF following in vivo microdialysis. Thus, EV^ISF^ isolations were performed via ultracentrifugation for all following experiments.

**Figure 3:**
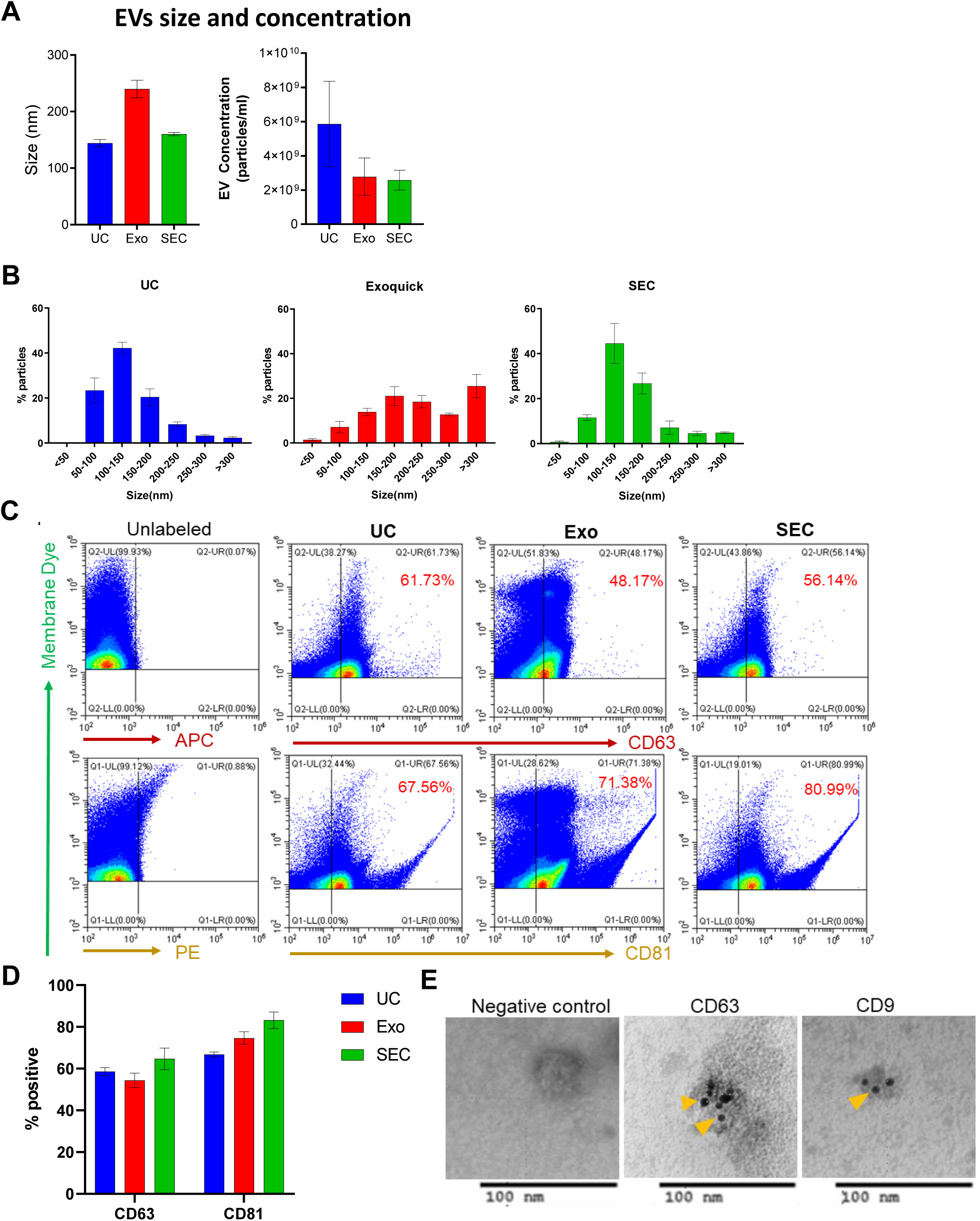
Characterization of EV^ISF^ isolated from ISF by multiple methods. EV^ISF^ were isolated from ISF samples (n=6) using three different methods; UC=Ultracentrifuge, Exo=ExoQuick and SEC= Size Exclusion Chromatography. **(A)** Nanoparticle tracking analysis (NTA) was performed to estimate the mean size and concentration (particles/ml) of EV^ISF^, isolated using UC (n=6), Exo (n=6) and SEC (n=3) methods. Graphs represent mean ± SEM. **(B)** Size distribution of particles was calculated from the NTA data and plotted as % particle for various sizes. **(C)** Flow cytometry analysis was performed to analyze the CD63+ and CD81+ EVs percentage in EV^ISF^ (n=3 each for method). EV^ISF^ were labeled with membrane labeling dye (CellBrite 488) and followed by CD63 (APC) and CD81 (PE) tagged antibodies. Dye positive EVs (representing membrane bound particles) on Y-axis were gated and analyzed for CD63/81 positive EVs. EV^ISF^ without antibody was used to set the gate for APC/PE positive EVs. Right shift in the APC/PE fluorescence showed the positive EVs. **(D)** The CD63+ and CD81+ EVs in EV^ISF^, analyzed by flow cytometry, isolated with different methods (n=3 each method) were plotted as mean ± SEM. **(E)** Immuno-gold labeling was performed to confirm the presence of CD63 and CD9 on EV isolated by UC method. Solid yellow arrows represent the presence of these markers on EVs surface.

### Age and Aβ pathology differentially modulate EV^ISF^ characteristics in APP/PS1 and WT mice

ISF was collected from 3-and 9-month-old WT and APP/PS1 mice for EV^ISF^ isolation and characterization. In accordance with the literature, no Aβ deposition was found in the hippocampus or cortex of APP/PS1 male and female mice at 3-months of age (Figure 4A-B) (45). By 9-months, Aβ deposition was present in both male and female APP/PS1 mice (Figure 4A-B). Female APP/PS1 mice had increased hippocampal Aβ deposition (p=0.0113), while amyloid plaque burden was less in males of the same age (Figure 4B) (45). In cortex, plaque pathology increased with age in both female (p<0.0001) and male APP/PS1 mice (p=0.0026), with amyloid plaque pathology more demonstrative in female than male mice (Figure 4B).

**Figure 4:**
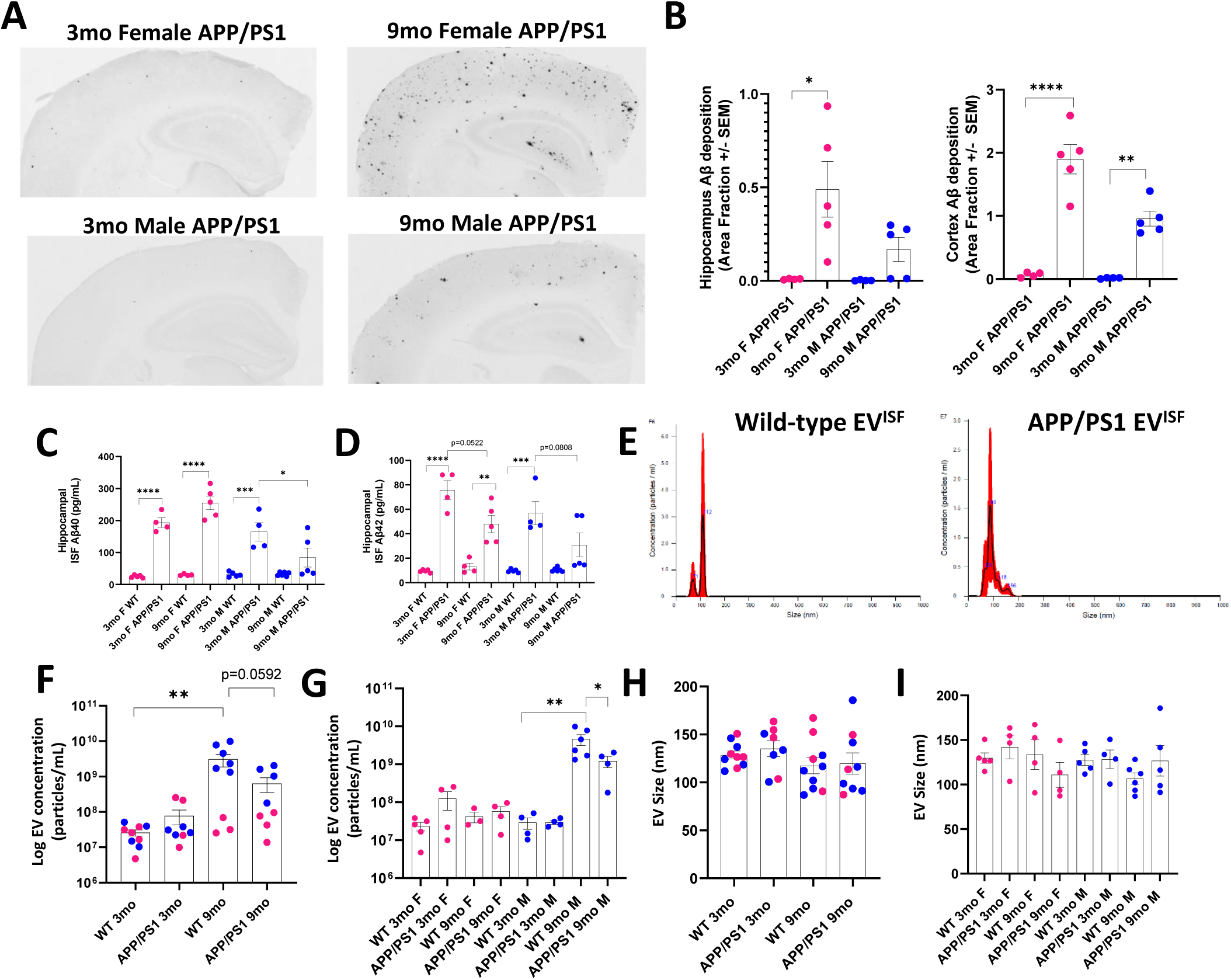
Age and Aβ pathology related changes in ISF Aβ, Aβ deposition, and EV^ISF^ characteristics in APP/PS1 and WT mice. **(A)** Representative images of Aβ deposition in 3mo and 9mo APP/PS1 mice, both sexes. **(B)** Aβ deposition in hippocampus and cortex was quantified and presented as Area Fraction. Plaque burden in hippocampus increased with age in female APP/PS1 mice (p=0.0113) but not in males. In cortex of APP/PS1, Aβ deposition increased in both females (p<0.0001) and males (p=0.0026) with age. **(C)** ELISAs for Aβ40 were performed on hippocampal ISF samples. ISF Aβ40 levels were higher in APP/PS1 compared to WT (3mo F WT vs 3mo F APP/PS1: p<0.0001, 9mo WT F vs 9mo F APP/PS1: p<0.0001, 3mo M WT vs 3mo M APP/PS1: p=0.0001) except for in 9mo WT vs 9mo APP/PS1 males. ISF Aβ40 levels decreased in 9mo APP/PS1 males (p=0.0449) but not females. **(D)** ELISAs for Aβ42 were performed on hippocampal ISF samples. ISF Aβ42 levels were increased in APP/PS1 mice compared to WT (3mo F WT vs 3mo F APP/PS1: p<0.0001, 9mo WT F vs 9mo F APP/PS1: p=0.0066, 3mo M WT vs 3mo M APP/PS1: p=0.0001) except for in 9mo WT vs 9mo APP/PS1 males. ISF Aβ42 levels had a trending decrease with plaque pathology in APP/PS1 in both males (p=0.0808) and females (p=0.0522). **(E)** NTA was performed on EV^ISF^. Representative size distribution and concentration of EV^ISF^ from WT and APP/PS1 3mo mice are shown. **(F)** EV^ISF^ concentration (particle number/mL) increased with age in WT mice (p=0.0098) but not APP/PS1. However, 9mo APP/PS1 EV^ISF^ concentration had a trending decrease compared to 9mo WT (p=0.0592). **(G)** EV^ISF^ concentration was separated by sex, age, and genotype. EV^ISF^ concentration increased with age in WT male mice (p=0.0028). EV^ISF^ concentration decreased with plaque pathology between 9mo WT and 9mo APP/PS1 males (p=0.0458) but not females. **(H)** EV^ISF^ size was the same across genotypes and age groups. **(I)** There was no difference in EV^ISF^ size across ages, sexes, and genotypes. p<0.05*, p<0.01**, p<0.001***, p<0.0001****. One-way ANOVA, multiple comparisons Tukey’s tests. N=3-6 per group

The ISF pool of Aβ is highly dynamic and reflects acute changes in the production and clearance of Aβ, leading to changes in Aβ aggregation and amyloid plaque formation (39). Therefore, we characterized how ISF Aβ levels change as a function of pathology, age, and sex in APP/PS1 and WT mice. While several studies explored how ISF Aβ levels changed relative to Aβ CSF or tissue burden, no study to date has investigated changes in ISF Aβ species relative to age, genotype, and sex (39). Due to transgenic overexpression, ISF Aβ40 levels were greater in APP/PS1 mice compared to WT. Interestingly, this effect was lost in 9-month-old APP/PS1 males compared to age-matched WT, where ISF Aβ40 levels decreased in APP/PS1 males (p=0.0449). Interestingly, we did not observe the same reduction in ISF Aβ40 in 9-month-old females, suggesting a sexual dimorphic response to ISF Aβ40 proteostasis and amyloid plaque formation (Figure 4C). ISF Aβ42 levels increased in APP/PS1 mice compared to WT with the exception of 9-month-old APP/PS1 males compared to 9-month WT males. ISF Aβ42 levels trended towards a decrease with plaque pathology in APP/PS1 for both males (p=0.0808) and females (p=0.0522) compared to age and sex-matched WT (Figure 4D). Together, these data demonstrate Aβ species-, sex-, age-, and pathology-dependent changes in ISF Aβ levels relative to Aβ aggregation.

EV^ISF^ from WT and APP/PS1 mice were characterized for size and concentration via NTA (Figure 4E-I). EV^ISF^ concentration increased with age in the WT (p=0.0098) but not in the APP/PS1 mice (Figure 4F). Interestingly, 9-month APP/PS1 mice trended towards a decrease in EV^ISF^ concentration compared to age-matched WT (p=0.0592; Figure 4F). When stratified by sex, EV^ISF^ concentration increased with age in WT males (p=0.0028). EV^ISF^ concentration decreased in 9-month-old APP/PS1 males compared to age-, sex-matched WT mice (p=0.0458; Figure 4G). This suggests a potential sex-specific change in EV^ISF^ uptake or release when amyloid plaques are present in APP/PS1 males (Figure 4G). EV^ISF^ size was unchanged with genotype, age, or sex (Figure 4H-I).

### Proteomics of total hippocampal EV^ISF^

LC-MS/MS proteomic results from total hippocampal EV^ISF^ revealed 436 proteins in EV^ISF^ present in at least 2 or more mice (Supplemental Table 1). DAVID analysis of all 436 proteins resulted in Gene Ontology (GO) terms for specific “Biological processes” which were grouped into related categories including proteostasis (32%), RNA processing (19%), immune function (10%), cell growth (6%), cellular trafficking (6%), glucose metabolism (6%), vascular health (6%), wound healing (4%), cellular excitability (3%), cellular structure (3%), apoptosis (2%), circadian rhythms (1%), DNA repair (1%), and lipid metabolism (1%) (Figure 5A). This confirms that the EV population isolated from the hippocampal ISF has shared characteristics of known EV functions and relationships (64–75). The top 10 GO terms for “Biological process”, “Molecular function”, KEGG pathways, and Canonical pathways were identified using DAVID and IPA analyses (Figure 5B). These top 10 results were clustered into categories related to proteostasis, inflammation/immune function, metabolism, and cellular structural integrity (Figure 5B). Further DAVID analysis of the EV^ISF^ proteomic data confirmed hippocampus as the top GO term “Tissue”, validating the efficacy of this technique to detect EV^ISF^ in a particular brain region (Figure 5C). DAVID analysis also verified extracellular exosomes as the top GO term for “Cellular component” in the EV^ISF^ dataset (Figure 5D). Comparison of the 436 proteins found in EV^ISF^ to the Top 100 ExoCarta proteins list (59) and a proteomic EV dataset from Choi et al. (58) revealed 49 shared proteins among the three groups (Figure 5E). Interestingly, 54 proteins were unique to EV^ISF^. EV^ISF^ also shared 29 proteins with EVs isolated from postmortem mouse brain tissue (60) and human brain tissue (15). This comparison revealed 168 proteins unique to hippocampal EV^ISF^ (Figure 5F). Assessment of the unique EV^ISF^ proteins from both Venn diagrams (Figures 5E-F) resulted in 45 proteins unique to EV^ISF^ only (See Supplemental Table 2). Thus, although hippocampal EV^ISF^ collected via in vivo microdialysis shared cargo with EVs isolated with other techniques, they are a unique pool of EVs with distinctive cargo not typically present in other EV datasets.

**Figure 5:**
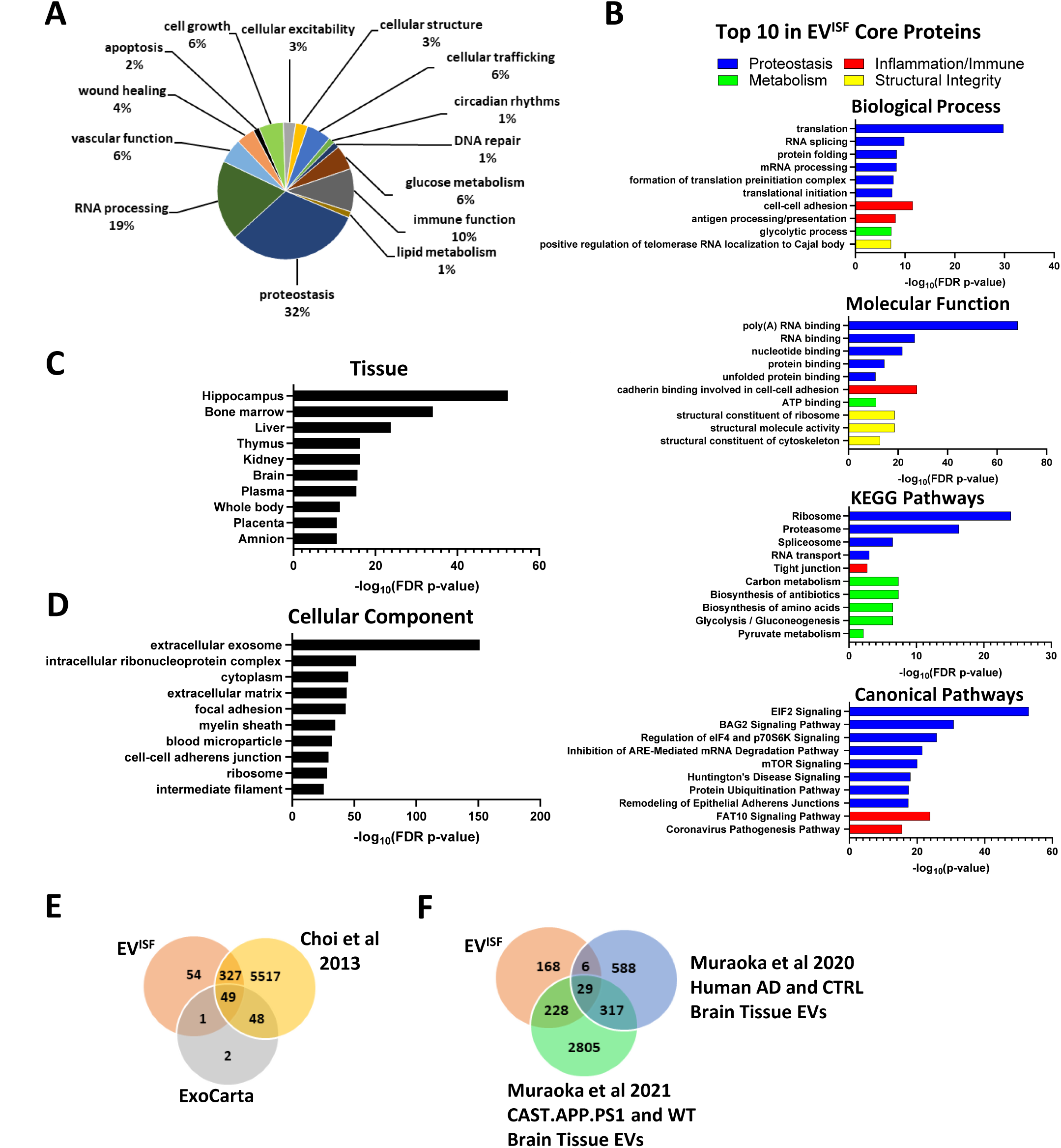
EV^ISF^ Proteomics: **(A)** Biological process groups for all EV^ISF^ proteins. **(B)** Top 10 Biological Processes, Molecular Functions, KEGG Pathways, and IPA Canonical Pathways for all EV^ISF^ proteins. Blue=Proteostasis, Red=Inflammation/Immune, Green=Metabolism, and Yellow=Structural Integrity. **(C)** DAVID analyses were performed on all the isolated EV^ISF^. The top UP_Tissue result was Hippocampus. **(D)** The top GOTERM_Cellular Component result was “extracellular exosome”. **(E)** Venn Diagram comparing EV proteins between the Choi et al 2013, ExoCarta Top 100, and EV^ISF^ datasets. **(F)** Venn Diagram comparing EV proteins between the Muraoka et al 2020, Muraoka et al 2021, and EV^ISF^ datasets.

### Age and Aβ pathology differentially alter EV^ISF^ protein concentration, protein diversity, and biological processes

EV^ISF^ protein concentration increased with age in both WT (p=0.0006) and APP/PS1 (p<0.0001; Figure 6A). However, the total number of identified proteins in the EV^ISF^ decreased with genotype, age, and Aβ deposition (Figure 6B). There were fewer proteins in 3-month APP/PS1 (p=0.0020), 9-month WT (p=0.0054), and 9-month APP/PS1 EV^ISF^ (p=0.0113) compared to 3-month WT (Figure 6B). The number of shared and unique EV^ISF^ proteins among all four groups were examined (Figure 6C). 168 proteins were shared among all groups, revealing novel brain EV markers that may be specific to the hippocampus. 3-month WT mice had the following 20 unique EV^ISF^ proteins: SNRPE, STRAP, PPP2CB, SRSF10, SARNP, PDIA3, PLEC, HSPA4L, FUBP1, CBX3, RPL37A, PSMD6, CCT4, RBMX, NME2, FH1, ELAVL1, RPS16, HSPA9, and PPIB. Interestingly, only two proteins were specific to 9-month WT EV^ISF^, TNNT3 and HMGB1. Surprisingly, no EV^ISF^ proteins were unique to 3-month-old APP/PS1 mice. Five proteins were unique to the 9-month-old APP/PS1 group: GLUD1, FBXL21, ALDH2, PGLYRP1, and NGP. Comparisons of the proteins across the four groups showed decreased protein diversity due to both genotype and age in APP/PS1 compared to WT (Figure 6D). There were fewer unique proteins (<u>Unique:</u> 3mo WT: 83; 3mo APP/PS1: 23) as well as an overall decrease in total protein number (<u>Total:</u> 3mo WT: 357; 3mo APP/PS1: 297) in APP/PS1 mice compared to WT at 3-months of age. Moreover, 9-month-old APP/PS1 mice had a further reduction in unique proteins compared to 3-month APP/PS1 (<u>Unique:</u> 3mo APP/PS1: 99; 9mo APP/PS1: 74) as well as a reduction in total protein number (<u>Total:</u> 3mo APP/PS1: 297; 9mo APP/PS1: 272). Additionally, 9-month-old APP/PS1 had fewer unique proteins compared to age-matched WT (<u>Unique:</u> 9mo WT: 92; 9mo APP/PS1: 19) as well as a lower number of total proteins (<u>Total:</u> 9mo WT: 345; 9mo APP/PS1: 272). The greatest drop in protein diversity was observed in 9-month APP/PS1 to 3-month WT (<u>Unique:</u> 3mo WT: 159; 9mo APP/PS1: 74), which was commensurate with the greatest decrease in total protein number (<u>Total:</u> 3mo WT: 357; 9mo APP/PS1: 272). Age-related changes in protein number were not as demonstrative, where 9-month WT had approximately the same variety of unique proteins compared to 3-month WT (<u>Unique:</u> 3mo WT: 86; 9mo WT: 74) as well as a similar number of total proteins (<u>Total:</u> 3mo WT: 357; 9mo WT: 345). Thus, although protein concentration increased with both age and pathology, APP/PS1 genotype and plaque pathology decreased protein diversity compared to age-matched controls.

**Figure 6:**
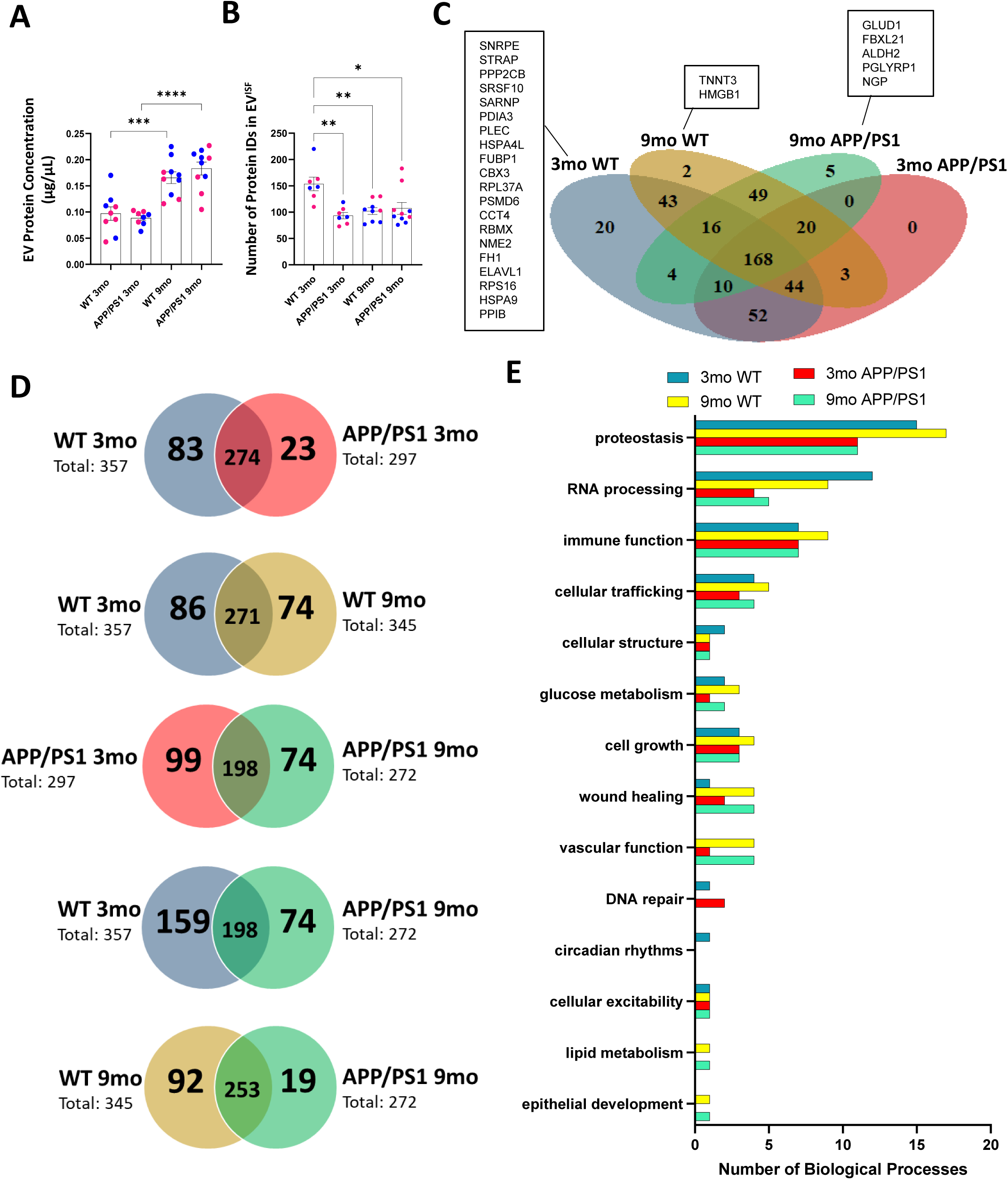
Protein concentration increases with age and pathology while protein diversity decreases in EV^ISF^. **(A)** EV^ISF^ protein concentration increased with age in both the WT (p=0.0006) and APP/PS1 mice (p<0.0001). **(B)** The number of unique protein IDs in EV^ISF^ decreased with genotype (p=0.0020), age (p=0.0054), and pathology (p=0.0113) compared to 3mo WT. **(C)** Venn Diagram reveals unique and shared EV^ISF^ proteins among all four groups: 3mo WT, 9mo WT, 3mo APP/PS1, and 9mo APP/PS1. **(D)** Venn diagrams of proteins unique to and shared between 3mo WT vs 3mo APP/PS1, 3mo WT vs 9mo WT, 3mo APP/PS1 vs 9mo APP/PS1, 3mo WT vs 9mo APP/PS1, and 9mo WT vs 9mo APP/PS1. Protein diversity decreased with genotype, age, and pathology. **(E)** DAVID analyses revealed changes in the number of biological processes within groups with age, genotype and pathology. One-way ANOVA, multiple comparisons Tukey’s tests p<0.05*, p<0.01**, p<0.001***, p<0.0001**** N=7-10 per group.

DAVID bioinformatics analyses were performed on EV^ISF^ proteins from 3-and 9-month-old WT and APP/PS1 mice. Using DAVID, Gene Ontology (GO) terms for “Biological process” within each group were determined, and results with-log(FDR-value) ≥ 1.3 were clustered into related categories including proteostasis, RNA processing, immune function, cellular trafficking, glucose metabolism, cell growth, wound healing, vascular function, DNA repair, circadian rhythms, cellular excitability, lipid metabolism, and epithelial development (Figure 6E). The number of biological processes that fell within each category were determined. Proteostasis, RNA processing, immune function, and cellular trafficking categories contained the greatest number of biological processes across all 4 groups. The number of proteostasis-related biological processes increased with age in WT compared to 3-or 9-month APP/PS1 groups. Both age and APP/PS1 genotype reduced the number of RNA processing-related biological processes in EV^ISF^. Interestingly, an increase in the number of immune function-related biological processes was observed in 9-month WT mice when compared to 3-month WT, 3-month APP/PS1, and 9-month APP/PS1 groups. Age-related increases in the number of wound healing-, vascular function-, lipid metabolism-, and epithelial development-related biological processes were observed in both APP/PS1 and WT mice. Conversely, the number of biological processes related to DNA repair mechanisms specifically increased in APP/PS1 at 3-months but decreased with age regardless of genotype.

### Bioinformatic analysis of EV^ISF^ proteins reveals differences in biological pathways associated with genotype, age, and plaque pathology

DAVID bioinformatics analyses were performed on EV^ISF^ proteins from 3-and 9-month-old WT and APP/PS1 mice. Using DAVID, Gene Ontology (GO) terms for “Biological process” within each group were determined, and results with-log(FDR-value) ≥ 1.3 were clustered into related categories as described previously (see Figure 6E). 3-month APP/PS1 EV^ISF^ had fewer total number of biological processes compared to 3-month WT (3mo WT: 49; 3mo APP/PS1: 36) (Figure 7A). Nevertheless, there was no difference in the percentage of proteostasis-related biological processes between 3-month WT and APP/PS1 (3mo WT: 31%; 3mo APP/PS1: 31%) (Figure 7A). However, in 3-month APP/PS1 EV^ISF^, the percentage of biological processes related to RNA processing decreased compared to WT (3mo WT: 25%; 3mo APP/PS1: 11%) (Figure 7A). This also corresponds with a decrease in protein diversity in 3-month-old APP/PS1 EV^ISF^ compared to age-matched WT (see Figures 6B and 6D). On the other hand, the percentage of immune function-related biological processes increased in 3-month-old APP/PS1 vs 3-month WT EV^ISF^ (3mo WT: 14%; 3mo APP/PS1: 19%) (Figure 7A). This suggests EV^ISF^ signal and/or propagate an early inflammatory response in 3-month APP/PS1 before the presence of plaque pathology is detectable. DAVID analysis of the 3-month WT and APP/PS1 EV^ISF^ revealed the top 10 GO terms for “Biological Process” and top 10 KEGG pathways while IPA revealed the top 10 canonical pathways. These results were grouped into the following categories: proteostasis, inflammation/immune, metabolism, or structural integrity (Figure 7B). In the top 10 “Biological Process” results, RNA splicing, mRNA processing, formation of translation preinitiation complex, and translational initiation, all processes related to proteostasis, were present in 3-month WT EV^ISF^ but not detected in 3-month APP/PS1 (Figure 7B). Conversely, the GO terms proteolysis involved in cellular protein catabolic process, positive regulation of protein localization to Cajal body, microtubule-based process, and intermediate filament organization were gained in the top 10 “Biological Process” results in 3-month APP/PS1 EV^ISF^ (Figure 7B). This suggests that genotype alters proteostasis and structural integrity processes. For the top 10 KEGG pathway results, RNA transport and pentose phosphate pathways were present in 3-month WT but not detected in 3-month APP/PS1 EV^ISF^ (Figure 7B). Nevertheless, phagosome and legionellosis KEGG pathways were gained in 3-month APP/PS1 (Figure 7B). IPA results revealed that in the top 10 canonical pathways of 3-month APP/PS1 EV^ISF^, mTOR signaling and protein ubiquitination pathway were absent but 14-3-3-mediated signaling and gluconeogenesis I pathways were gained compared to 3-month WT (Figure 7B). These results demonstrate that EV^ISF^ can detect and/or propagate proteostasis, inflammation/immune, and metabolic changes taking place between WT and APP/PS1 as early as 3-months of age.

**Figure 7:**
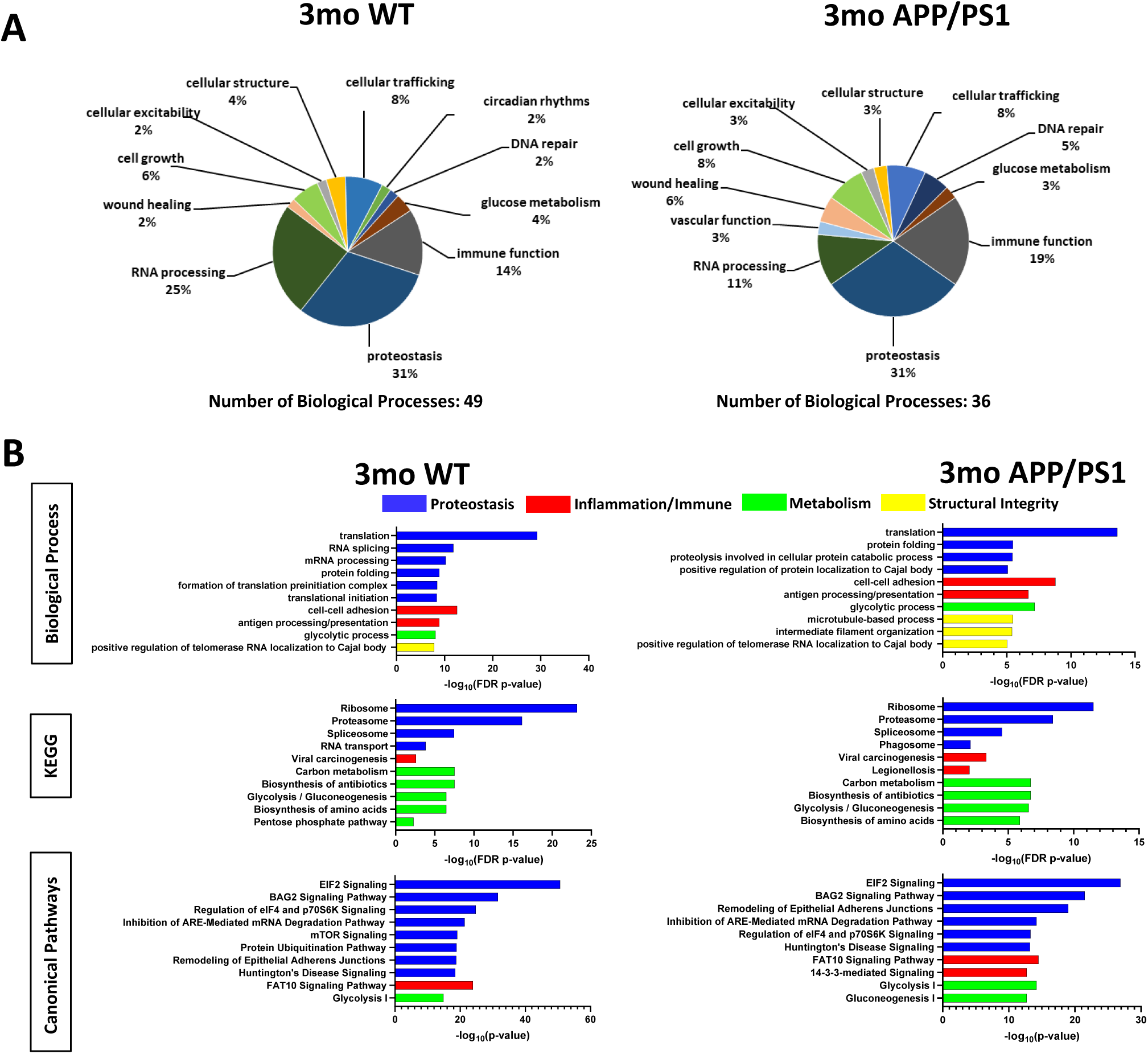
EV^ISF^ reveal early changes in proteostasis and inflammation in APP/PS1. DAVID analyses were performed on 3mo WT and 3mo APP/PS1 EV^ISF^. **(A)** Biological process results were grouped into related categories. **(B)** Top 10 Biological Processes, KEGG Pathways, and IPA Canonical Pathways for 3mo WT and 3mo APP/PS1 EV^ISF^ proteins, respectively. Blue=Proteostasis, Red=Inflammation/Immune, Green=Metabolism, and Yellow=Structural Integrity.

In 9-month APP/PS1 EV^ISF^, the total number of biological processes was lower compared to age-matched WT (9mo WT: 59; 9mo APP/PS1: 44) (Figure 8A). However, there was little difference in the percentage of biological process categories between 9-month WT and APP/PS1. For example, the percent of proteostasis-and RNA processing-related biological processes were similar between 9-month APP/PS1 and 9-month WT EV^ISF^ (<u>Proteostasis:</u> 9mo WT: 29%; 9mo APP/PS1: 25%; <u>RNA</u> <u>processing:</u> 9mo WT: 15%; 9mo APP/PS1: 12%). This potentially underscores the important age-component of AD. For the top 10 “Biological Process” results, the GO terms regulation of translational initiation, formation of translation preinitiation complex, positive regulation of RNA polymerase II transcriptional preinitiation complex assembly, cell-cell adhesion, and positive regulation of telomerase RNA localization to Cajal body were lost from 9-month WT to 9-month APP/PS1 EV^ISF^ (Figure 8B). However, 9-month APP/PS1 EV^ISF^ gained protein folding, response to peptide hormone, acute-phase response, microtubule-based process, and muscle contraction in the top 10 “Biological Processes” compared to 9-month WT (Figure 8B). In the top 10 KEGG pathways, 9-month APP/PS1 lost spliceosome and gained systemic lupus erythematosus compared to age-matched WT EV^ISF^ (Figure 8B). The top 10 IPA canonical pathways revealed that regulation of eIF4 and p70S6K signaling, mTOR signaling, protein ubiquitination pathway, Huntington’s disease signaling, and FAT10 signaling pathway were lost in 9-month APP/PS1 EV^ISF^ vs 9-month WT (Figure 8B). In contrast, 9-month APP/PS1 EV^ISF^ gained the canonical pathways glucocorticoid receptor signaling, LXR/RXR Activation, acute phase response signaling, glycolysis I, and actin cytoskeleton signaling in the top 10 results vs age-matched WT (Figure 8B), suggesting changes in immunometabolism in APP/PS1 mice compared to age-matched controls.

**Figure 8:**
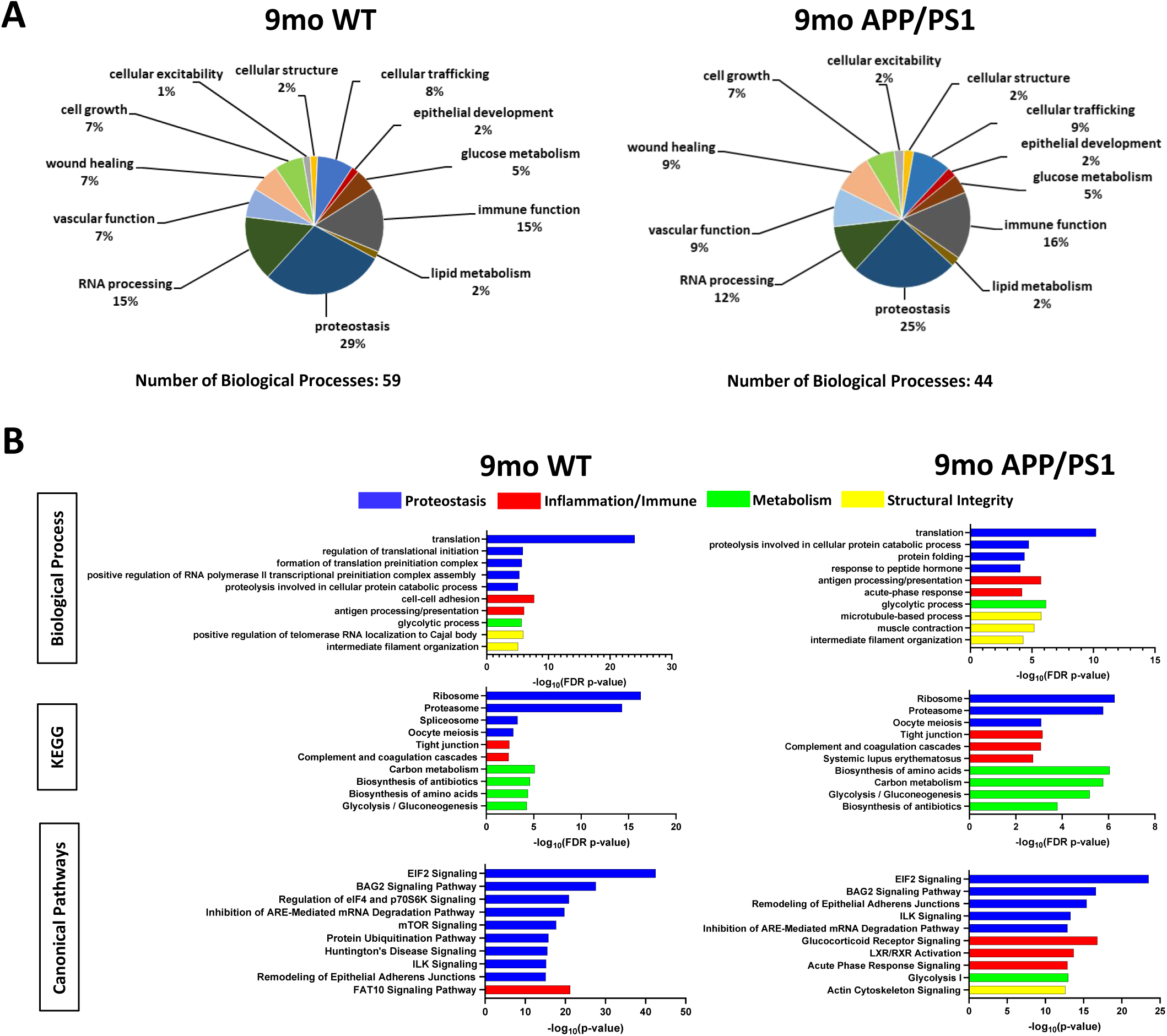
EV^ISF^ reveal changes in proteostasis and inflammation with Aβ deposition. DAVID analyses were performed on 9mo WT and 9mo APP/PS1 EV^ISF^ proteomic results. **(A)** Biological process results were grouped into related categories. **(B)** Top 10 Biological Processes, KEGG Pathways, and IPA Canonical Pathways for 9mo WT and 9mo APP/PS1 EV^ISF^, respectively. Blue=Proteostasis, Red=Inflammation/Immune, Green=Metabolism, and Yellow=Structural Integrity.

### Percentages of cell type-related proteins in EV^ISF^ are modulated by and correlate with ISF Aβ levels in both sexes

Next, we explored whether EV^ISF^ proteins originated from neurons, astrocytes, oligodendrocytes, microglia, and/or endothelial cells using the Barres Lab brainrnaseq.org database (49) (Figure 9A). The percentages of proteins from each cell type were calculated then represented as a percentage of the total pool for each of the four groups (Figure 9B). We found that cell type-specific changes in the EV^ISF^ proteome were modulated by genotype, age, pathology, and sex. The percent of endothelial cell-related proteins decreased as a function of genotype (p=0.0222), age (p=0.0110), and Aβ pathology (p=0.0626) in both male and female mice compared to 3-month WT (Figure 9C). Conversely, the percentage of oligodendrocyte-related proteins did not change across groups (Figure 9C). 9-month-old APP/PS1 mice trended towards a decrease (p=0.0814) in EV^ISF^ neuron-related proteins compared to 9-month WT (Figure 9D). The percentage of neuron-related proteins in EV^ISF^ positively correlated with ISF Aβ40 (p=0.0326, r=0.5050) and ISF Aβ42 (p=0.0061, r=0.6195) levels in APP/PS1 mice (Figure 9D). Interestingly, the percentage of neuron-related proteins in EV^ISF^ did not correlate with Aβ deposition (Figure 9D). This suggests that EV^ISF^ neuronal proteins correlate with ISF Aβ levels before amyloid plaque formation. While astrocyte-related proteins in EV^ISF^ were unchanged between groups (Figure 9E), the percentage of astrocyte-related proteins in EV^ISF^ positively correlated (p=0.0168, r=0.5549) with ISF Aβ40 levels. There was no correlation between astrocyte-EV^ISF^ proteins and ISF Aβ42 or Aβ deposition (Figure 9E). There was no change in the percentage of microglia-related proteins in EV^ISF^ across groups (Figure 9F). However, microglia-EV^ISF^ proteins negatively correlated with ISF Aβ40 (p<0.0001, r=-0.7931) and ISF Aβ42 (p=0.0214, r=-0.5376) levels but not with Aβ deposition (Figure 10F).

**Figure 9:**
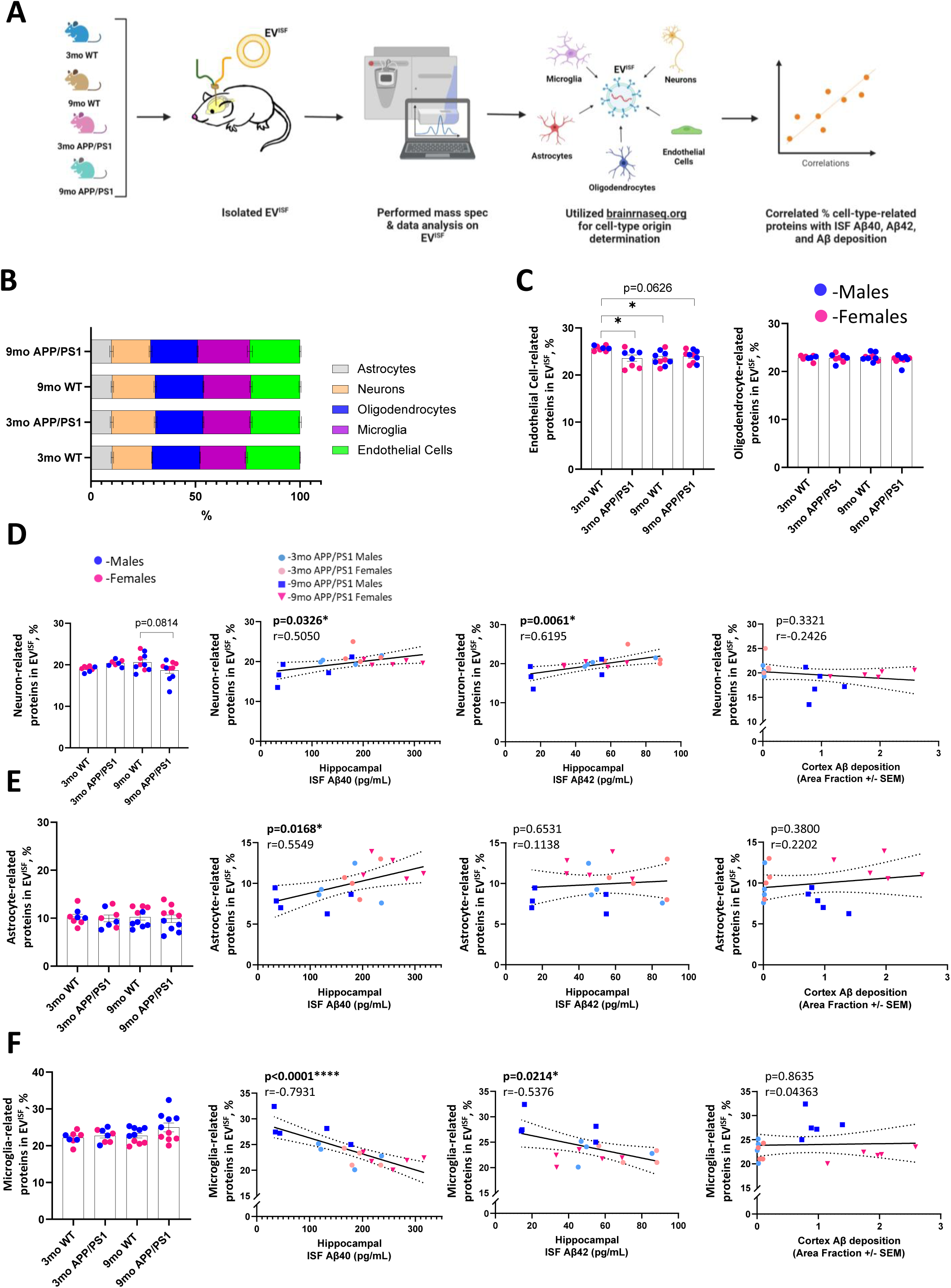
Aβ pathology and age modulate cell type-related EV^ISF^ proteins. **(A)** Illustration of cell-sources of EV^ISF^ proteins. **(B)** EV^ISF^ proteins were searched in the brainrnaseq.org database (49) and were grouped according to their cell-source(s) per mouse. Percentages represent the average of each cell type-source per group. **(C)** In EV^ISF^, endothelial cell-related proteins decreased with APP/PS1 genotype (p=0.0222) and age (p=0.0110). They trended down with pathology compared to 3mo WT (p=0.0626). There was no difference among groups for oligodendrocyte-related proteins in EV^ISF^. **(D)** Neuron-related proteins trended down with pathology compared to 3mo APP/PS1. Percentage of neuron-related proteins correlated with ISF Aβ40 and Aβ42 levels but not Aβ deposition in APP/PS1 mice. **(E)** The percentages of astrocyte-related proteins in EV^ISF^ were unchanged across groups. However, they positively correlated with ISF Aβ40. **(F)** The percent of microglia-related proteins in EV^ISF^ did not change among groups. However, they did negatively correlate with ISF Aβ40 and Aβ42 levels. One-way ANOVAs with Tukey multiple comparisons test. p<0.05* Endothelial: N=8-10/group, Astrocyte: N=8-10/group, Oligodendrocyte: N=8-10/group, Neuron: N=7-10/group. Microglia=8-10/group. Correlation: Pearson r and two-tailed p-value: p<0.05*, p<0.0001****. N=18

**Figure 10:**
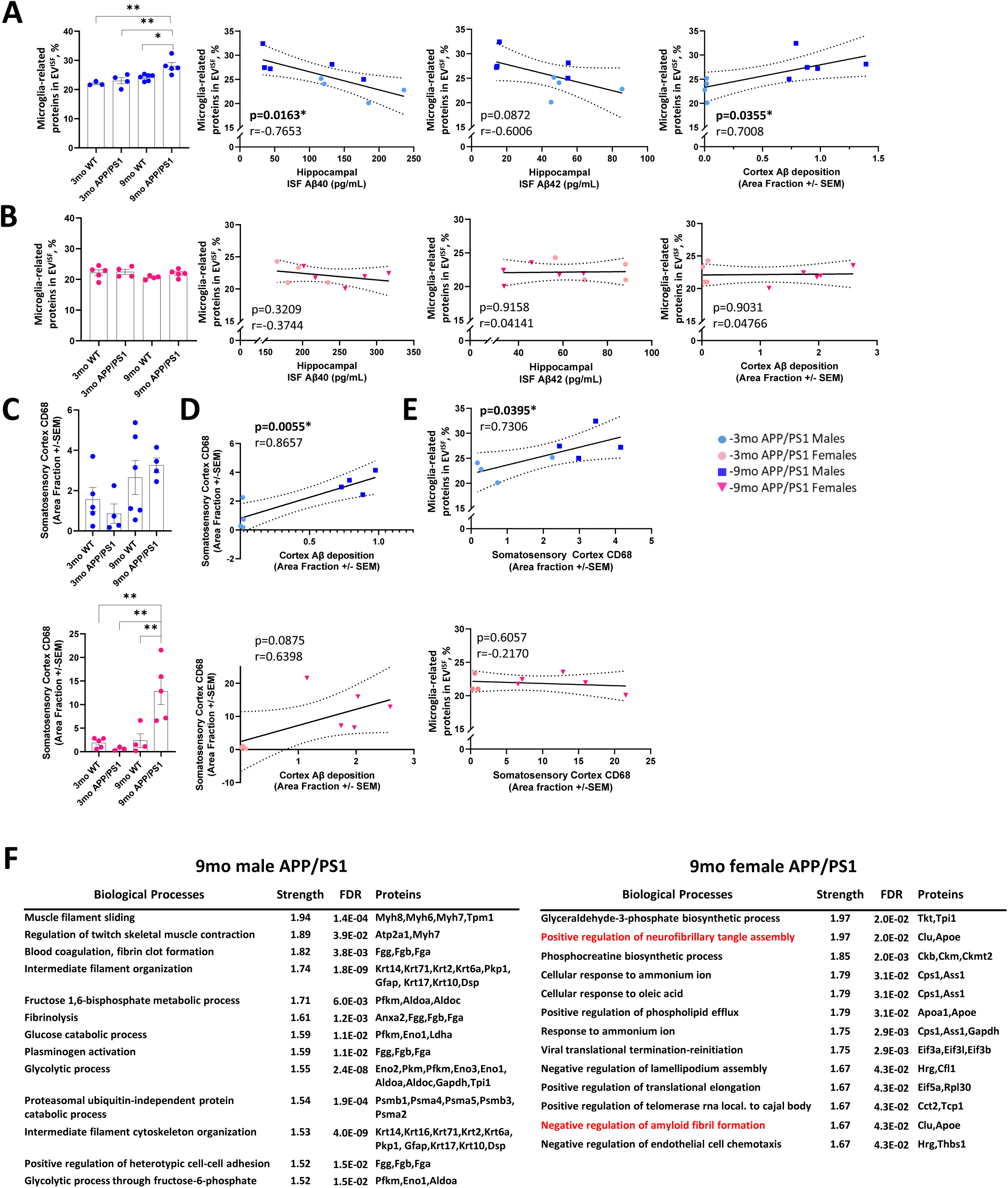
Microglia-derived EV^ISF^ proteins correlate with ISF Aβ, Aβ deposition, and microgliosis, sex-dependently, in APP/PS1. **(A)** EV^ISF^ proteins were searched in the brainrnaseq.org database (49) and were grouped as described above. Percentages represent the average of each cell type source per group. When separated by sex, the percentage of EV^ISF^ microglia-related proteins in males increased with age and pathology (Males: 3mo WT vs 9mo APP/PS1: p=0.0036, 3mo APP/PS1 vs 9mo APP/PS1: p=0.0074, 9mo WT vs 9mo APP/PS1: p=0.0208). There was a negative correlation with ISF Aβ40 levels and a positive correlation with Aβ deposition. There was a trending negative correlation with ISF Aβ42. **(B)** In contrast, there was no difference in the percentage of microglia-related proteins among groups in females. Also, microglia-proteins in EVs did not correlate with ISF Aβ40, ISF Aβ42, or Aβ deposition. **(C)** Only 9mo APP/PS1 females had increased CD68+ levels. **(D)** However, CD68+ microglia levels in APP/PS1 male mice correlated with Aβ deposition while females only had a positive trend. **(E)** The percentage of microglia-related proteins in EV^ISF^ positively correlated with CD68+ microglia staining only in male APP/PS1. **(F)** Separate STRING analyses of 9-month APP/PS1 males and females revealed biological processes unique to each sex. Further, neurofibrillary tangle and amyloid-related biological processes in the top 20 results in females only. One-way ANOVAs with Tukey multiple comparison tests: p<0.05*, p<0.01**, p<0.001***, p<0.0001****, N=3-10/group. Correlation: Pearson r and p-value: p<0.05*

### Microglia-related EV^ISF^ proteins correlate with activated microglia (CD68^+^) in APP/PS1 in a sex-dependent manner

When our results were further stratified by sex, we observed sex-dependent effects of Aβ pathology on microglia-related proteins in the EV^ISF^ population. Specifically, the percentage of microglia-related proteins increased with the presence of plaque pathology in males but not females (Figure 10A). The percent of microglia-related proteins negatively correlated with ISF Aβ40 levels (p=0.0163, r=-0.7653) only in APP/PS1 males. There was also a trend towards a negative correlation between microglia-related proteins and ISF Aβ42 levels only in APP/PS1 males (p=0.0872, r=-0.6006; Figure 10A). However, the percentage of microglia-related proteins in EV^ISF^ positively correlated with Aβ deposition in male APP/PS1 (p=0.0355, r=0.7008; Figure 10A). In female mice, the percentage of microglia-related proteins in EV^ISF^ did not change due to genotype, age, or pathology (Figure 10B). Moreover, microglia-related EV^ISF^ proteins did not correlate with ISF Aβ40, ISF Aβ42 levels, or Aβ deposition in APP/PS1 females (Figure 10B).

In addition to amyloid plaque burden, we also assessed the relationship between microglia-related proteins in EV^ISF^ and disease-associated microglia (DAM), CD68^+^ microglia. Consistent with previous results (76,77), we observed an increase in CD68^+^ staining in 9-month-old APP/PS1 mice compared to 3-month APP/PS1, 3-month WT, or 9-month WT mice (Figure 10C). There was no difference across groups for CD68^+^ microglia staining in males (Figure 10C). However, female 9-month APP/PS1 had more CD68^+^ microglia than 3-month WT (p=0.0027), 3-month APP/PS1 (p=0.0035), and 9-month WT (p=0.0059) mice (Figure 10C). CD68^+^ staining positively correlated with Aβ deposition in males (p=0.0055, r=0.8657) and trended towards a positive correlation with Aβ deposition in females (p=0.0875, r=0.06398; Figure 10D). Interestingly, microglia-related proteins in EV^ISF^ positively correlated with CD68^+^ microglia staining in APP/PS1 males (p=0.0395, r=0.7306; Figure 10E). Surprisingly, microglia-related EV^ISF^ proteins and CD68^+^ staining did not correlate in female APP/PS1 (Figure 10E). This demonstrates that when levels of microglia activation in the brain increase in response to plaque pathology, levels of microglia-related proteins in the EV^ISF^ of 9-month-old APP/PS1 males increase commensurately. Finally, due to the differences seen between male and female microglia-related proteins, STRING analyses were performed on the EV^ISF^ proteins from 9-month APP/PS1 male and female mice, separately (Figure 10F). The 9-month male and female APP/PS1 mice demonstrated unique and shared biological processes in their top 20 results. Muscle filament sliding, regulation of twitch skeletal muscle contraction, blood coagulation, fibrin clot formation, intermediate filament organization, fructose 1,6-bisphosphate metabolic process, fibrinolysis, glucose catabolic process, plasminogen activation, glycolytic process, proteasomal ubiquitin-independent protein catabolic process, intermediate filament cytoskeleton organization, positive regulation of heterotypic cell-cell adhesion, and glycolytic process through fructose-6-phosphate were biological processes unique to the male, 9-month, APP/PS1 EV^ISF^ (Figure 10F). On the other hand, the biological processes glyceraldehyde-3-phosphate biosynthetic process, positive regulation of neurofibrillary tangle assembly, phosphocreatine biosynthetic process, cellular response to ammonium ion, cellular response to oleic acid, positive regulation of phospholipid efflux, response to ammonium ion, viral translational termination-reinitiation, negative regulation of lamellipodium assembly, positive regulation of translational elongation, positive regulation of telomerase RNA localization to Cajal body, negative regulation of amyloid fibril formation, and negative regulation of endothelial cell chemotaxis were unique to the 9-month, female, APP/PS1 EV^ISF^ (Figure 11F). Interestingly, only female, 9-month APP/PS1 demonstrated two AD-associated biological processes, 1) positive regulation of neurofibrillary tangle assembly and 2) negative regulation of amyloid fibril formation. These results were due to the presence of the AD-related proteins apolipoprotein E (APOE) and clusterin (CLU) in the 9-month, APP/PS1, female EV^ISF^ proteome (Figure 10F). Finally, both sexes shared the following biological processes in their top 20 results: peptidyl-cysteine s-trans-nitrosylation, neutrophil aggregation, methylglyoxal biosynthetic process, telomerase holoenzyme complex assembly, regulation of vacuole fusion, non-autophagic, peptidyl-cysteine s-nitrosylation, and protein refolding. Therefore, the sex-dependent, microglial protein differences demonstrated in EV^ISF^ are driven by an AD-related phenotype.

**Figure 11:**
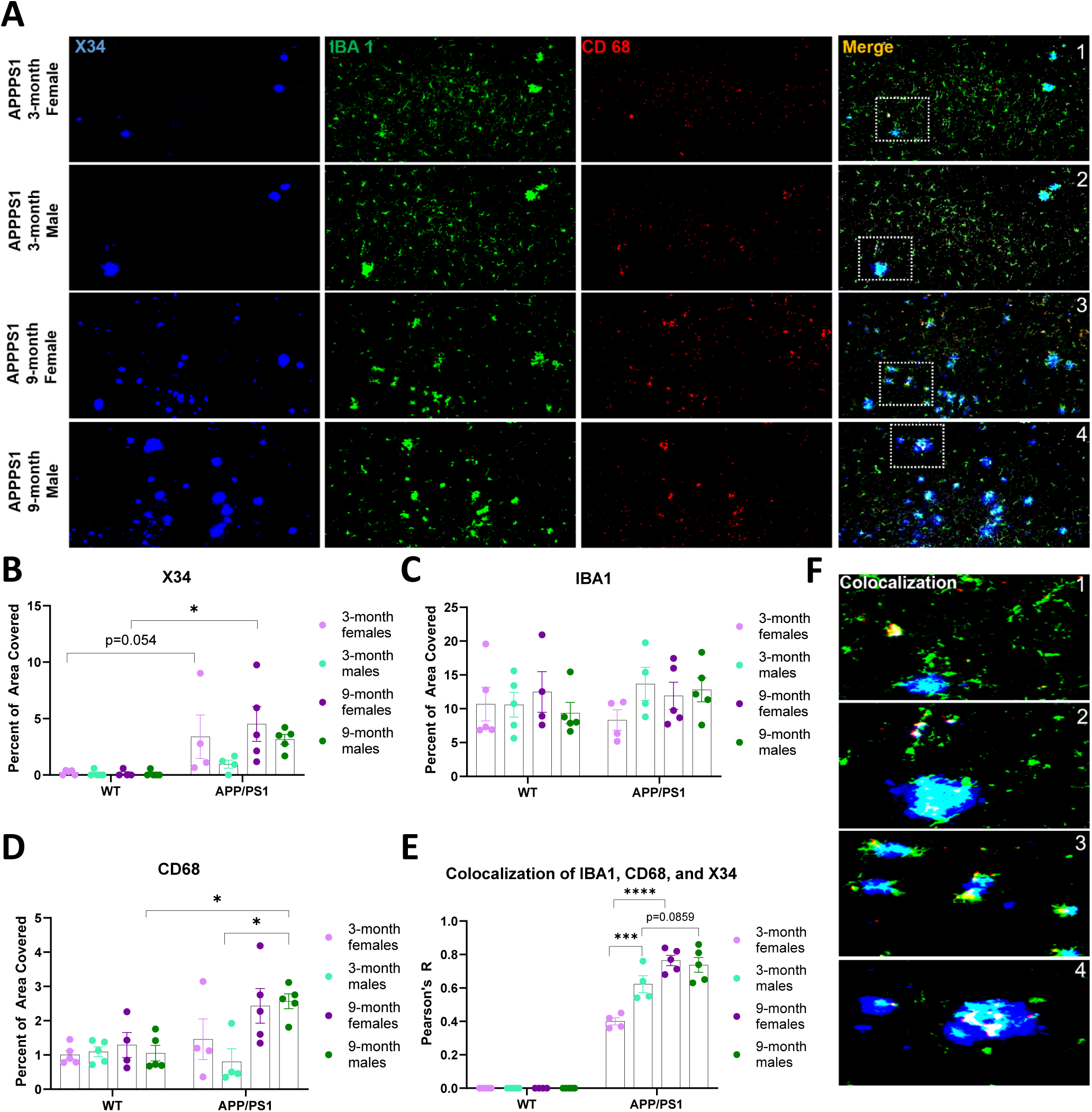
Male and female microglia differentially associate with amyloid plaques. **(A)** Representative images of the anterior cortex taken from 3-and 9-month-old APP/PS1 mice. **(B)** Plaque coverage was quantified using X34. A main effect of genotype (p=0.0014) driven by female APP/PS1 mice was found. Post hoc analyses revealed 9-month female APP/PS1 mice had greater plaque coverage than WT (p=0.0313) with a similar trend at 3-months. **(C)** There were no statistical differences in macrophage coverage identified using IBA1. **(D)** Phagocytic activity was measured using CD68. There were main effects in genotype (p=0.0064) and age (p=0.0042) and a genotype x age interaction (p=0.0148). Post hoc analyses revealed that CD68 area coverage increased with age and plaque pathology in APP/PS1 male mice (p=0.0218). Further, 9mo APP/PS1 males had greater CD68 levels than 9mo WT (p=0.0460). **(E)** Colocalization analyses were performed to determine the involvement of microglia phagocytic activity in X34 deposition. There were main effects of genotype (p<0.0001), age (p<0.0001), and sex (p=0.0259). Three-way interaction revealed a main effect of genotype x age (p<0.0001), genotype x sex (p=0.0259), sex x age (p=0.0041), and sex x age x genotype (p=0.0041). Post hoc analyses revealed that 3-month APP/PS1 males had higher rates of colocalization than females (p=0.0002). Further, colocalization increased with age and plaque pathology in APP/PS1 females (p<0.0001) with a similar trend in APP/PS1 males. **(F)** Representative images of colocalization. Images are numbered 1-4 in relation to their original image in (A). Mixed-effects analyses with Tukey’s multiple comparison tests: p<0.05*, p<0.01**, p<0.001***, p<0.0001****, N=4-5/group.

### Activated microglia in APP/PS1 mice differentially engage with amyloid plaques, sex-dependently

Since EV^ISF^ revealed immunometabolic-related changes in their proteomes when Aβ pathology was present, microglia and their association with amyloid plaques were assessed in 3-month and 9-month old WT and APP/PS1 mice. Additionally, both homeostatic and activated, DAM microglia were examined across groups. Anterior cortex sections were co-stained for homeostatic microglia (IBA1), DAM microglia (CD68), and amyloid plaques (X34) (Figure 11A). APP/PS1 mice had greater plaque deposition when compared to WT (p=0.0014; Figure 11B). This genotype effect was driven by APP/PS1 females. 3-month APP/PS1 females had a trending increase in plaque deposition compared to age-matched WT (p<0.054) while 9-month APP/PS1 females had more amyloid plaques than WT mice (p=0.0313; Figure 11B). Plaque burden in APP/PS1 male mice was unchanged when compared to WT mice or APP/PS1 females (Figure 11B). IBA1 staining was unchanged across groups (Figure 9C). On the other hand, CD68 staining revealed main effects in genotype (p=0.0064) and age (p=0.0042) as well as a genotype x age interaction (p=0.0148). CD68 increased in male APP/PS1 from 3-month to 9-month (p=0.0218; Figure 11D). 9-month APP/PS1 male mice also exhibited greater CD68 coverage when compared to 9-month WT males (p=0.0460; Figure 11D). However, CD68 did not change between 3-month and 9-months in APP/PS1 females (Figure 11D). Thus, activated, DAM microglia but not homeostatic microglia were increased only in male APP/PS1 mice when Aβ pathology was present. Finally, the colocalization of X34, IBA1, and CD68 was assessed across groups to identify whether activated microglia were differentially recruited to amyloid plaques. Colocalization revealed main effects with genotype (p<0.0001), age (p<0.0001), and sex (p=0.0259). Further, there were main effects of genotype x age (p<0.0001), genotype x sex (p=0.0259), sex x age (p=0.0041), and sex x age x genotype (p=0.0041) interactions. Interestingly, APP/PS1 males demonstrated greater colocalization than females at 3-months (p=0.0002; Figure 9E-F). However, colocalization increased in APP/PS1 female mice with age and plaque pathology (p<0.0001) while males had a trending increase (p=0.0859; Figure 11E). Thus, activated microglia differentially colocalize with amyloid plaques in a sex-dependent manner in APP/PS1 mice, where male mice show a greater number of plaque associated microglia early in life that is related to less amyloid pathology at late stages and a more robust EV^ISF^ microglial proteome signature.

## Discussion

We developed a novel and innovative method for examining the role of brain EVs in AD. This approach leveraged in vivo microdialysis as a way to isolate small EVs from the brain’s ISF (EV^ISF^) in unanesthetized, freely moving mice. Not only were we able to circumvent potential limitations associated with current approaches, but we also made novel insights into the relationship between hippocampal EV^ISF^ and the progression of AD. The EV field needs new approaches to isolate EVs from the brain’s extracellular space without collecting ILVs or other extracellular non-vesicles (61). Historically, in vivo microdialysis has successfully collected proteins and metabolites directly from the brain’s ISF (35, 39, 41, 51, 52) yielding novel insights into how this dynamic pool is regulated. To date, Aβ_1-x_ (35, 37, 39), Aβ38 (78), Aβ40 (37, 39, 42, 78), Aβ42 (37, 39, 78), tau (41,52), ApoE (63), glucose (35,42), lactate (35,37), and ethanol (42) are all detectable in the ISF of mouse models of AD-related pathology. For the first time, we collected EV^ISF^ via in vivo microdialysis and analyzed their size, concentration, and proteome. Moreover, we investigated changes in EV^ISF^ relative to age, sex, Aβ pathology, and neuroinflammation. Using this approach, we found that while the total EV^ISF^ concentration and protein concentration in EV^ISF^ increases with pathology, the diversity of the EV^ISF^ proteome decreased with Aβ aggregation, specifically in proteins originating from neurons and endothelial cells. Interestingly, the increase in the EV^ISF^ cargo proteins was largely due to microglial-derived proteins. Changes in the microglial EV^ISF^ proteome were sexually dimorphic and associated with a differential response of plaque associated microglia. We found that the female APP/PS1 mice have more amyloid plaques, less plaque associated microglia, and a less robust-and diverse-EV^ISF^ microglial proteome. Pathway analysis on the EV^ISF^ proteome from female APP/PS1 mice showed two pro-Alzheimer’s related pathways driven by two AD associated risk proteins, APOE and CLU, that are known to promote Aβ aggregation (79,80). These proteins were not detected in EV^ISF^ from the male APP/PS1 mice. Together, we not only developed a novel method of collecting EVs from the hippocampal ISF, but also demonstrated a sexually dimorphic response of microglia to Aβ pathology that is related to the microglial EV^ISF^ proteome. These observations not only suggest a role for EVs in microglia-amyloid plaque interactions, but also has implications for EV-based biomarker development which historically focuses on neuronal-or astrocyte-derived EVs to stage or diagnosis AD.

Our novel in vivo microdialysis approach is important because it avoids caveats from existing methods. To date, brain-EVs are isolated from postmortem, not live, brain tissue. While this includes the study of brain-EVs from humans, rodent models, and non-human primates, postmortem analyses have caveats. The postmortem interval can alter RNA expression and integrity (81), shift brain metabolism (82), and accelerate protein degradation (83). Thus, postmortem tissue changes may affect brain-derived EVs by causing artifactual changes in the EV cargo that would be absent if isolated from a live brain. Several brain tissue-derived EV techniques use harsher isolation protocols, resulting in potential intracellular and non-vesicular contamination (20, 21, 23, 25, 26, 84). Since ILVs retain many of the same markers as secreted EVs, it is hard to delineate whether an EV pool is found intracellularly or extracellularly with current techniques. In vivo microdialysis collection of EV^ISF^ circumvents these caveats. Our approach also allows us to explore EVs from a specific brain region vulnerable to AD pathology during amyloid plaque formation (85). Sampling brain-derived EVs from biofluids like blood or CSF represents a heterogeneous pool of brain-derived EVs that may or may not have originated from regions impacted by AD-related pathology. Additionally, samples are traditionally examined distal to the brain in humans, monkeys, and rodents which relies on intact transport and clearance mechanisms, which are dysfunctional in AD and other CNS diseases. Despite advances in the EV field, it is still difficult to isolate and compare EVs from specific brain regions even in postmortem tissue (21, 27, 86, 87). Because of the limited number of human or nonhuman primate samples, often only one brain region from a few samples can be studied at a time. While more brain regions and a greater sample size can be employed using rodent models, size constraints limit region-specific isolation of brain-EVs with current methods. An entire brain hemisphere, a whole brain, or pooled mouse brains are generally required for EV isolation (22-24, 88, 89). Thus, differences in EVs from different brain regions are missed. This is imperative for studying CNS disease since not all brain regions are uniformly impacted by disease or undergo pathological changes at the same rate. In contrast, changes in peptides, proteins, or metabolites due to pathology, age, or pharmacological intervention can be measured in the ISF of specific brain regions using in vivo microdialysis (35, 36, 46, 90). We have now expanded this approach to measure changes in EVs as it relates to AD-related pathology.

Given what is well established in the EV field, several of our study’s key findings were unexpected. First, our approach revealed sex-dependent changes in EV^ISF^ concentration due to age not observed in previous studies. EV^ISF^ concentration increased as male mice aged but not females (Figure 4F-G). This is in contrast to a plasma-EV study that found EV concentration decreased with age (91). These changes could reflect age-or pathology-related changes in clearance mechanisms that plague most CNS diseases, including AD. We also found protein concentration within hippocampal EV^ISF^ increased with age in WT mice (Figure 6A) while protein diversity remained the same (Figures 6B and 6D). In contrast with hippocampal EV^ISF^, the contents of plasma-EVs decreased with age (92). Interestingly, the number of proteostasis-related biological processes increased as the mice aged in our EV^ISF^ (Figure 6E). These results agree with studies demonstrating that aging affects proteostasis via alterations in protein synthesis, degradation, chaperones, the ubiquitin-proteasome system, and autophagy (93). Next, we explored how EV^ISF^ protein concentration and diversity changed with amyloid pathology. Here, we observed an age-dependent increase in EV^ISF^ protein concentration, but a striking decrease in protein diversity in the 9-month APP/PS1 mice (Figures 6A-B and 6D). We hypothesized that since changes in protein synthesis, trafficking, degradation, and clearance are all affected in AD, changes in EV^ISF^ proteome reflect this. Indeed, the number of proteostasis-related biological processes as well as the number of proteostasis-related pathways decreased with APP/PS1 genotype and Aβ deposition compared to age-matched WT (Figures 6E, 7B, and 8B). These results agree with the literature demonstrating that proteostasis is dysregulated in aging and AD (94–96). Thus, AD-related changes in proteostasis appear to be modulating protein diversity, biological processes, and pathways within hippocampal EV^ISF^ population.

Next, we demonstrate that the EV^ISF^ proteome changes in a cell type-specific manner *in vivo* despite a global increase in EV^ISF^ protein concentration with age. While percentage of oligodendrocyte-and astrocyte-related proteins in EV^ISF^ are unchanged with genotype, age, or Aβ deposition (Figure 9C), we did see that endothelial-, neuronal-, and microglial-associated cargo proteins were affected by age, sex, or genotype. For example, endothelial cell-related proteins in hippocampal EV^ISF^ decreased with genotype and age and had a trending decrease as mice developed amyloid plaques (Figure 9C). However, changes in endothelial-related proteins in EV^ISF^ did not correlate with changes in ISF Aβ40, ISF Aβ42, or amyloid pathology. In humans, endothelial cell-EVs increased with disease in plasma from AD patients (97). This difference could be due to the fact that vascular-derived EVs are secreted into blood vessel lumen, not brain parenchyma, as the disease develops. Therefore, changes in protein abundance would be reflected in the plasma, not ISF EV pool. Conversely, it could suggest decreases in the vascular-EV proteome may be more important to pursue as an AD biomarker when staging disease progression. While astrocyte-related proteins in EV^ISF^ were unchanged across groups (Figure 9E), we did find a positive correlation between astrocyte-related proteins in EV^ISF^ and ISF Aβ40 levels. It is also interesting that astrocyte-related proteins in EV^ISF^ correlate with ISF Aβ40, but not ISF Aβ42 or amyloid pathology. ISF Aβ40 is the more abundant species of Aβ and less aggregatory prone compared to Aβ42 (98–101), but also highly regulated by changes in excitatory neurotransmission (39, 40, 102). We hypothesize the relationship between ISF Aβ40 and astrocyte-related proteins in EV^ISF^ reflects astrocyte-neuron cooperation in the healthy brain, which is lost with the onset of Aβ aggregation. In fact, when examining the astrocyte-derived EV^ISF^ proteome in 9-month APP/PS1 mice using STRING, there is a shift towards pathways involved in Aβ/tau aggregation, cellular metabolism, and inflammation (data not shown). While this suggests that the EV pool of astrocyte-related proteins is stable across genotype, sex, and age, exploring how specific proteins change in astrocyte-derived proteins holds promise as AD biomarkers in agreement with previous studies (103). In our EV^ISF^ population, neuron-related proteins had a trending decrease with Aβ deposition (Figure 9D) but positive correlations with ISF Aβ40 and ISF Aβ42 levels. Similar to the findings with the EV^ISF^ astrocyte-derived proteome, this suggests that neuronal activity drives EV^ISF^ release in a manner similar to Aβ_1-x_ (37, 38, 40). It also suggests that as Aβ40 and Aβ42 levels drop in the ISF, which occurs during Aβ aggregation (104), neuron-derived proteins in EV^ISF^ also decrease. This is consistent with previous studies that found neuronal-related markers in brain tissue-EVs were decreased in AD patients compared to controls (15,86). However, many studies still focus on changes in neuronal-derived EVs or exosomes as a biomarker of AD pathology. If less neuronal-derived proteins are synthesized, packaged, or trafficked in EVs in regions undergoing degeneration, then neuronal-derived EVs may not be the most attractive biomarker in AD. It would be interesting to explore the relationship between neuronal-derived EVs and AD further.

The most interesting results from our study were related to EV^ISF^ microglia-related proteins (Figure 9–10). First, prior to the formation of amyloid plaques, EV^ISF^ from 3-month APP/PS1 mice had a greater number of inflammation-related pathways compared to 3-month WT (Figure 7B). Following Aβ deposition, there are more immunometabolic-related canonical pathways in EV^ISF^ compared to controls (Figure 8B), highlighting the role of neuroinflammation in AD as well as other neurodegenerative diseases (105–107). Next, increases in the EV^ISF^ microglia-related proteins were negatively correlated with ISF Aβ40 and ISF Aβ42, suggesting that an increase in EV^ISF^ microglial-proteins is associated with drops in both ISF Aβ40 and Aβ42, or Aβ aggregation. When separated by sex, EV^ISF^ microglia-related proteins increased with Aβ deposition in male APP/PS1 mice and correlated with ISF Aβ40, ISF Aβ42, and most interestingly, cortical Aβ deposition in male mice (Figures 10A). We demonstrate that as early as 3-months of age when Aβ starts to aggregate, CD68+ microglia colocalize more strongly with amyloid plaques in male mice compared to females. This is despite more total CD68+ microglia in female mice (106–109). We also see that EV^ISF^ microglia-related proteins are positively correlated with amyloid plaques, but only in male mice, which also have less amyloid pathology than females. Comparing the total EV^ISF^ proteome at 9-months, pathway analysis revealed two AD-related pathways related to neurofibrillary tangle assembly and amyloid fibril formation are driven by the AD-related proteins and pro-Aβ aggregatory proteins, APOE and CLU, but only in female mice. Our studies demonstrate a sexually dimorphic response of microglia to amyloid plaques which may offer one explanation to why there is a differential risk for males and females in AD. Moreover, the EV^ISF^ microglial proteome differs by sex, shedding light on the sex-dependent role for microglia proteins and EVs in AD.

Given the strong relationship between EV^ISF^ microglia-related proteins and Aβ, this suggests that microglial-derived EVs could be an attractive, early biomarker of Aβ-related changes in the brain. Previous studies demonstrate that microglial-derived EVs can transport pathological tau and Aβ (86, 110, 111). Microglia can uptake EVs from the extracellular space, serving as a brain clearance mechanism for Aβ (112). Brain-EVs isolated from post-mortem AD models demonstrate that DAM microglia can excessively secrete EVs in the brain in the presence of AD-related pathology (113) and brain tissue-EVs express an enrichment of DAM proteins (60). To date, few studies have explored sexually dimorphic response of microglia to AD-related pathology and whether microglial derived-EVs are involved, which may offer one explanation to why there is a differential risk for males and females in AD. Our data provides mechanistic support for epidemiological studies that females are at greater risk for AD compared to males. Thus, our technique for hippocampal EV^ISF^ collection reveals previously undescribed changes in cell type-specific proteins with age, sex, genotype, and Aβ deposition. Moreover, it suggests that microglial-derived EVs hold promise for staging the progression of AD. Further examination of brain-EVs in vivo provides a unique opportunity to better understand the role brain-derived EVs, particularly microglial-derived EVs, play in AD as well as their role as potential biomarkers.

In conclusion, we developed and validated a novel approach using in vivo microdialysis to isolate sEVs from the ISF of the live brain to better understand the role of brain-derived EVs in AD. Our work builds upon the current EV and AD literature to capture how the dynamic, in vivo pool of EV^ISF^ changes with AD pathogenesis in a brain region vulnerable to disease. EV^ISF^ are altered with age. They reveal proteostasis changes that are modulated by the presence of Aβ deposition. More inflammatory-and immune-related pathways appear in EV^ISF^ with APP/PS1 genotype and plaque pathology compared to WT. Microglia-related proteins are sex-dependently increased in male 9-month APP/PS1 EV^ISF^. Thus, microglial contents of EV^ISF^ may be moderating a beneficial immunometabolic response that could be driving the sex differences seen not only in APP/PS1 mice but also in humans with AD (107,114). Studying EV^ISF^ will further elucidate the roles of EVs, microglia, and sex-differences in AD pathology.

## ACKNOWLEDGEMENTS

We would like to thank Dr. David Holtzman for generous gifts of Aβ antibodies.

## ETHICS APPROVAL

All animal procedures and experiments were performed under guidelines approved by the Institutional Animal Care and Use Committee (IACUC) at Wake Forest School of Medicine.

## AVAILABILITY OF DATA AND MATERIALS

The datasets used, produced, and/or analyzed during the current study are available from the corresponding author on reasonable request. Of note, all data generated or analyzed during this study are included in this published article. All proteomic datasets will be open access.

## AUTHOR CONTRIBUTIONS

SLM and GD conceived of the study. SLM, GD, and MCP contributed to study design. MCP, SDK, YS, AK, SS, SCG, SV, MA, CMC, JAS, and JL performed experiments. MCP, SDK, and SCG performed data analysis and data interpretation. MCP, SLM, and GD wrote the manuscript. All authors discussed the results and commented on the manuscript.

## COMPETING INTERESTS

SLM served as a consultant for Denali Therapeutics. Conflicts of interest: None

## FUNDING

We would like to acknowledge the following grants: Wake Forest Alzheimer’s Disease Research Center pilot and R01AG061805 (GD, SLM), P30AG072947 (SLM), K01AG050719 (SLM), R01AG068330 (SLM), BrightFocus Foundation A20201775S (SLM), Averill Foundation (SLM), F31AG071119 (MP), and Wake Forest Baptist Comprehensive Cancer Center Proteomics and Metabolomics Shared Resource, supported by the National Cancer Institute’s Cancer Center Support Grant award number P30CA012197.

## Notes

### Competing Interest Statement

The authors have declared no competing interest.

